# The sequences near Chi sites allow the RecBCD pathway to avoid genomic rearrangements

**DOI:** 10.1101/351395

**Authors:** Chastity Li, Claudia Danilowicz, Tommy F. Tashjian, Veronica G. Godoy, Chantal Prévost, Mara Prentiss

## Abstract

Bacterial recombinational repair is initiated by RecBCD, which creates a 3′ single-stranded DNA (ssDNA) tail on each side of a double strand break (DSB). Each tail terminates in a Chi site sequence that is usually distant from the break. Once an ssDNA-RecA filament forms on a tail, the tail searches for homologous double-stranded DNA (dsDNA) to use as template for DSB repair. Here we show that the nucleoprotein filaments rarely trigger sufficient synthesis to form an irreversible repair unless a long strand exchange product forms at the 3′ end of the filament. Our experimental data and modeling suggest that terminating both filaments with Chi sites allows recombinational repair to strongly suppress fatal genomic rearrangements resulting from mistakenly joining different copies of a repeated sequence after a DSB has occurred within a repeat. Taken together our evidence highlights cellular safe fail mechanisms that bacteria use to avoid potentially lethal situations.

## Introduction

While eukaryotes use complex strategies (Ryu et al., 2016, Amaral et al., 2017) to avoid dangerous rearrangements that can result when repeated sequences interfere with double strand break (DSB) repair (Bao et al., 2015, Ryu et al., 2016, Amaral et al., 2017), bacterial strategies have remained mysterious. Understanding the mechanisms for rejecting major rearrangements in bacterial genomes may provide better predictions of possible rearrangements. Furthermore, knowledge of the role of Chi sites in the DSB repair may influence the efficiency of gene targeting (Dabert and Smith, 1997).

When a DSB occurs in bacteria, it can be repaired using RecA-mediated homologous recombination following the well-known RecBCD pathway (Figure 1a) (Symington, 2014, Mawer and Leach, 2014, Azeroglu et al., 2016, Kowalczykowski, 2015, Smith, 2012, Smith, 1991). RecBCD degrades or resects each end of the broken double-stranded DNAs (dsDNA) until it recognizes a Chi site. Chi sites are ~8 bp DNA sequences that alter the function of RecBCD to create two 3′ ssDNA tails that terminate in Chi sites (Symington, 2014, Mawer and Leach, 2014, Azeroglu et al., 2016, Kowalczykowski, 2015, Smith, 2012, Smith, 1991) (Figure 1ai, ii). RecA then binds to the ssDNA tails, creating two ssDNA-RecA filaments with Chi sites at their 3′ ends. Those ssDNA-RecA filaments then search for homologous regions in the dsDNA.

**Figure 1.**
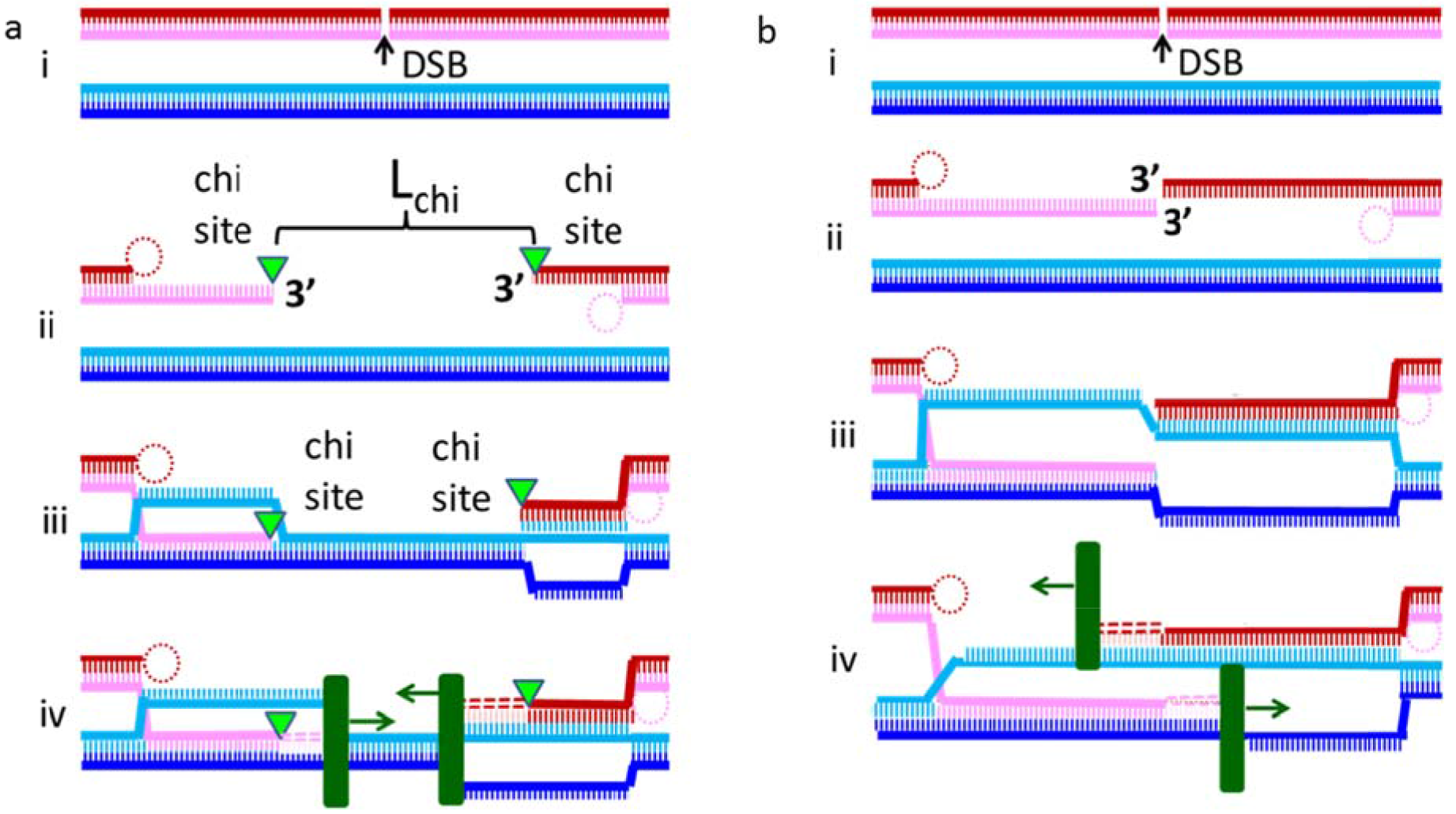
Model of the RecBCD-dependent and hypothetical DSB ends repair pathways. **a** Schematic of the RecBCD dependent DSB repair pathway. i. The dsDNA sequences are identical, but the dsDNA molecule with red and pink strands contains a double strand break indicated by the black arrow and labeled DSB. ii. RecBCD creates ssDNA with Chi sites (indicated by green triangles) at their 3′ ends, while the complementary strands are degraded or looped (dotted circles) creating an L_Chi_ bp gap. iii. RecA mediated strand exchange creates heteroduplex products that reach the 3′ ends of the filaments. iv. DNA polymerase (dark green rectangles) extends both initiating ssDNAs by copying the complementary strands beginning at the 3′ ends of an initiating strand in a RecA filament. This process is only irreversible once there are no unmatched bases between the initiating filament and the complementary strand. **b** Schematic for a hypothetical DSB break repair mechanism that is similar to (a) except the 3′ ends of the DSB form the 3′ ends of the searching filament and RecBCD does not participate. We compare results from this mechanism to results from the RecBCD pathway to determine whether removing L_Chi_ between the 3′ ends of the filament and positioning Chi sites at the 3′ ends of the searching ssDNA filaments allows the RecBCD pathway to suppress genomic rearrangement due to pairing between different copies of long repeated sequences.

To determine whether a region of dsDNA is homologous to the initiating strand, ssDNA-RecA filaments attempt strand exchange. Strand exchange establishes Watson-Crick pairing between the initiating strand and one of the strands in the dsDNA. The first sequence matching test attempts to establish base pairing between approximately 8 nt (Howard-Flanders et al., 1984, Danilowicz et al., 2015, Qi et al., 2015, Bazemore et al., 1997, Yang et al., 2015, Hsieh et al., 1992). Evidently, formation of the 3-strand heteroduplex product is most favorable if the heteroduplex is sequence matched. If at least 7 of the 8 bp match, RecA promotes formation of a metastable 8 bp heteroduplex product pairing bases in the initiating strand with bases in the complementary strand (Howard-Flanders et al., 1984, Danilowicz et al., 2015, Qi et al., 2015, Bazemore et al., 1997, Yang et al., 2015, Hsieh et al., 1992). Strand exchange can then extend the heteroduplex product in a 5′ to 3′ direction with respect to the initiating ssDNA (Mawer and Leach, 2014, Cox, 2007, Gupta et al., 1998) (Figure 1aiii). The stability of sequence matched strand exchange products increases strongly as the product length (L_prod_) increases from 8 to 20 bp (Hsieh et al., 1992, Danilowicz et al., 2015, Danilowicz et al., 2017, Qi et al., 2015), and *in vivo* results suggest that DNA repair is extraordinarily rare unless L_prod_ > 20 bp (Lovett et al., 2002, Watt et al., 1985, Shen and Huang, 1986).

In the presence of ATP hydrolysis, heteroduplex stability *in vitro* increases only slightly as L_prod_ extends from 20 to 75 bp (Danilowicz et al., 2017). If L_prod_ > 80 bp the nucleoprotein filament separates from the recombination complex (van der Heijden et al., 2008); however, even if L_prod_ > 80 bp, strand exchange products remain reversible (Rosselli and Stasiak, 1990, Danilowicz et al., 2017) unless two complete dsDNA strands are formed. In order to form two complete dsDNA strands, the bases removed by RecBCD (L_Chi_) must be replaced. That replacement is achieved by DNA synthesis that begins at the terminal 3′ OH on each initiating strand and uses the complementary strand as template (Figure 1aiv) (Li et al., 2009, Liu et al., 2011).

Figure 1b shows a hypothetical alternate pathway for DSB repair, in which the 3′ ends of the DSB form the 3′ ends of the searching filaments, and the process is otherwise identical to the RecBCD pathway (Wilkinson et al., 2016, Singleton et al., 2004, Kowalczykowski, 2000, Dillingham and Kowalczykowski, 2008) In this work, we will compare the genomic rearrangement that would be produced by this hypothetical pathway with the outcome of the RecBCD pathway to highlight the advantages conferred by the following two features: 1. removing L_Chi_ bases flanking the DSB, and 2. ensuring that the searching ssDNA strands have Chi sites at their 3′ ends. We will show that if repair follows the pathway shown in Figure 1a, these two features combined with the sequence distributions within bacterial genomes reduce or eliminate genomic rearrangements that would otherwise plague DSB repair.

## Results

### Long repeats are prevalent in bacterial genomes

Figure 2a, b illustrates that repeated sequences capable of forming a stable heteroduplex (L_prod_ > ~20 bp) (Hsieh et al., 1992, Danilowicz et al., 2015, Bazemore et al., 1997, Qi et al., 2015) are particularly prevalent in the *E. coli* O157 genome (gray lines in Figure 2a, b). In contrast, such repeats are rare in sequences consisting of randomly chosen bases (random sequences with the same length as the *E. coli* O157 genome, ~5 Mbp), as illustrated by the black lines in Figure 2a, b.

**Figure 2.**
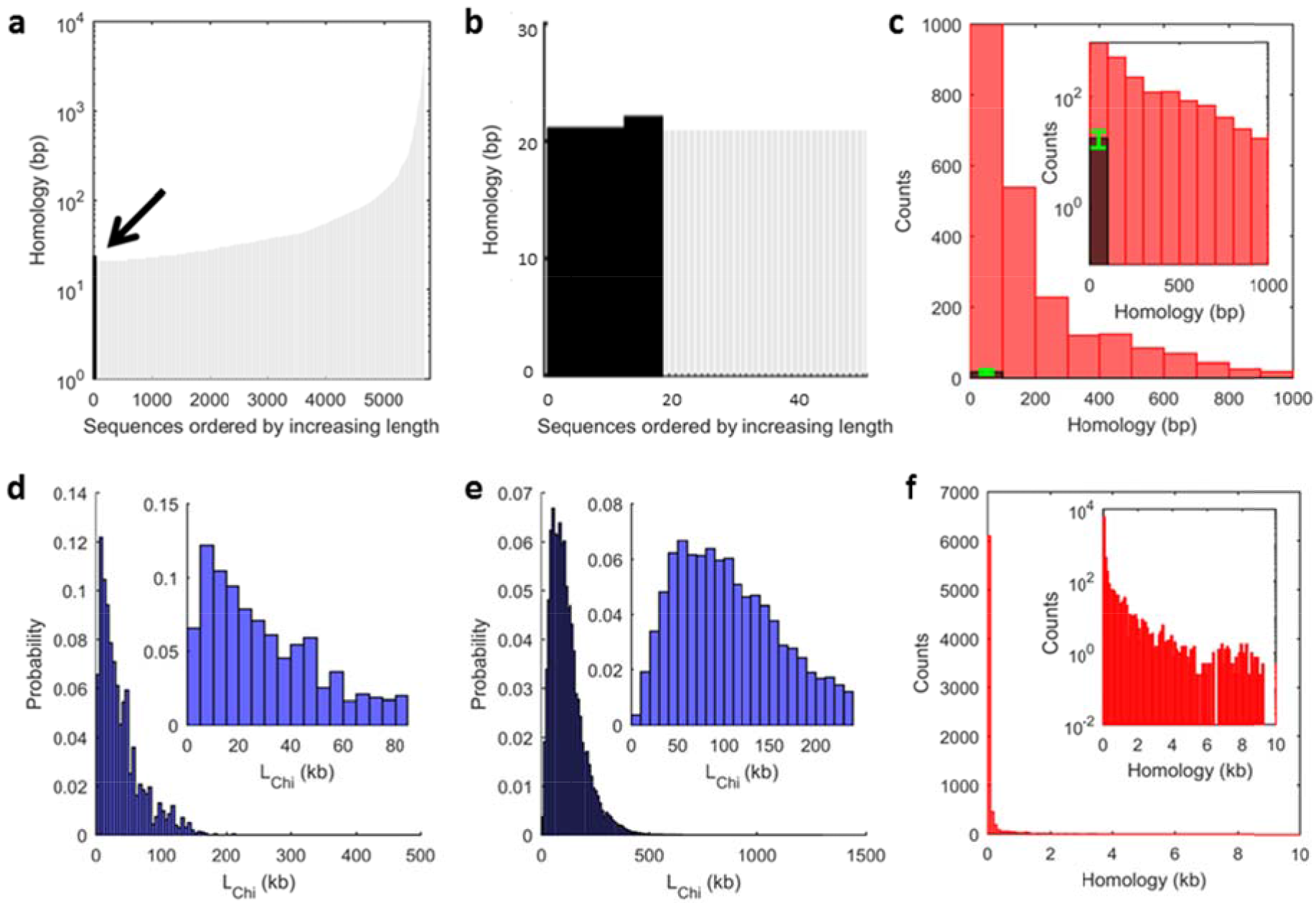
Prevalence of long repeats in bacterial genomes suggests rearrangements would be likely without RecBCD intervention. **a** Every repeated sequence in *E. coli* O157 with a length N_repeat_ > 20 is indicated by a gray line. The height of the line corresponds to the length of the repeat. The black line indicated by the arrow shows a typical result for a sequence of the same length whose bases were randomly chosen. **b** Same as (**a**) but with an expanded x-axis. For the 100 random sequences (about 5 Mb long) considered, the minimum and maximum number of repeats was 4 and 30, respectively. Mean = 17.6, mode = 18, and the longest repeat was 25 bp long. **c** The red bars in the histogram represents repeats between 20 < N_repeat_ < 1000 bp averaged over the four *E. coli* genomes using a 100 bp bin width. The dark bar shows the average of the results obtained for the 100 random sequences (about 5 Mb long). The green error bar shows the standard deviation for the 100 random sequences. Inset: same data using a logarithmic y-axis. **d** Probability distribution for L_Chi_ averaged over four *E. coli* genomes assuming 100 % Chi site recognition and using a 5 kb bin width. Inset: same data with an expanded x-axis. e Same as (**d**) but assuming 30 % Chi site recognition and using a 10 kb bin width. **f** Histogram of N_repeat_ averaged for the four *E. coli* genomes using a 100 bp bin width. The 10 kb maximum x-axis value corresponds to the bin width in (**e**). Inset: same as (**f**) but with a logarithmic y-axis showing that 8 % of repeats have lengths > 300 bp, and 4 % have lengths > 1 kb, whereas no repeat extends more than 9.5 kb.

The rarity of long repeats in random sequences of the same length as an *E. coli* genome is also illustrated by the dark gray bar clearly seen in the inset of Figure 2c, in which the histogram shows the number of repeated sequences of N_repeat_ (length of a repeated sequence occurring anywhere in the genome) that are longer than 20 bp and shorter than 1000 bp. The bar represents averages over 100 random sequences with lengths of ~5 Mbp. The green error bar shows the standard deviation. The data confirm that a homology test of 25-30 bp would be sufficient to prevent genomic rearrangements if bacterial sequences consisted of randomly selected bases (Vlassakis et al., 2013), since repeats shorter than ~25 bp rarely form irreversible products *in vivo* (Lovett et al., 2002, Watt et al., 1985, Shen and Huang, 1986).

Figure 2c also suggests that substantial genomic rearrangements are likely to occur if irreversible recombination products were to form between a 20-30 bp repeats anywhere in the genome. Though the method that we used to find long repeated sequences only finds exact repeats, long repeated regions containing some mismatches appear in the graph as several shorter exact repeats. We find that those exactly repeated shorter regions are almost never separated by more than one single base.

*In vivo* results indicate that the probability of recombining DNA increases exponentially as the homologous region in the recombining DNA strand extends from N = 20 to N = 75, where N = 75 is more than 100x more probable than N = 50 (Lovett et al., 2002, Watt et al., 1985). Remarkably, recombination increases only slightly as N increases from 75 to ~ 300 bp. It has been speculated that *in vivo* several parallel sequence-matched interactions with L_prod_ < 75 bp separated by ~200 bp may enhance discrimination against N_repea_t <~200-300 bp (Prentiss et al., 2015). Studies in *E. coli* suggest that RecA-dependent genomic rearrangements between directly repeated sequences in plasmids is improbable unless the repeat length is at least ~300 bp, though RecA independent rearrangements between shorter repeats do occur (Bi and Liu, 1994).

*In vivo* results mix the discrimination provided by RecA alone with the discrimination provided by other factors, and we note that not all *in vivo* recombination follows the RecBCD pathway. In the following, we will demonstrate how the RecBCD pathway reduces the probability that a DSB creates one searching filament that includes a region of a repeat with 75 < N < 300 bp at its 3′ end and eliminates the possibility that the *3′* ends of both filaments will include more than 20 bases that originate from the same repeat.

### Removing L_Chi_ bases by RecBCD promotes genomic stability

Without considering the detailed statistical distribution of Chi sites with respect to repeats, some advantages of the RecBCD pathway can be appreciated by considering a case in which a DSB occurs in the middle of a long repeated sequence. In the hypothetical DSB repair mechanism illustrated in Figure 1b, a DSB occurring within a repeated sequence will create two searching filaments whose 3′ ends terminate in regions of the repeated sequence that flanked the DSB. Genomic rearrangement will result if the two searching filaments pair with both sides of a different copy of the repeated sequence flanking the break.

In contrast, Figure 2 indicates that in the RecBCD pathway, which is illustrated in Figure 1a, the repeated sequence that flanked the DSB is likely to lie within the L_C_hi bases removed by RecBCD. In particular, Figure 2d indicates that the space between adjacent Chi sites on opposite strands is typically > 10 kb. Furthermore, Figure 2e indicates that since 30 % Chi site recognition is observed *in vivo* (Cockram et al., 2015, Taylor and Smith, 1992) an L_C_hi distribution that peaks at ~ 50 kb would be created.

Importantly, Figure 2f shows a histogram of the repeats averaged over four *E. coli* genomes. The maximum x-axis value in Figure 2f corresponds to the bin width size in Figure 2e. Thus, all of the repeats in the considered *E.coli* genomes have lengths that are smaller than 99.6 % of the L_Chi_ since the height of the first bin in Figure 2f is 0.4 %. This simple comparison of the maximum repeat length in the four *E. coli* genomes to the distribution of Lchi values makes it plausible that the removal of the L_chi_ bases surrounding a DSB could strongly suppress genomic rearrangement due to the two searching filaments pairing with both sides of a different copy of the repeated sequence flanking the break.

### Homology determines whether DNA synthesis stabilizes repairs

Other advantages of the RecBCD pathway emerge from more complex considerations that include detailed examination of bacterial sequences and experimental studies that determine what regions of the initiating ssDNA can lead to the DNA synthesis required for irreversible strand exchange and repair of the DSB. Previous work suggested that extension of the initiating strand by Pol IV may stabilize D-loops prior to re-establishment of a DNA polymerase III-dependent replication (Lovett, 2006), and that even in eukaryotic cells, translesion polymerases may aid DSB repair by stabilizing strand invasion intermediates (Lovett, 2006). This is consistent with new work indicating that most Pol IV molecules carry out DNA synthesis outside replisomes (Henrikus et al., 2018).

In these experiments, we study DNA synthesis by *E. coli* DNA Polymerase IV (Pol IV) as well as by the large fragment of *Bacillus subtilis* DNA polymerase I (LF-Bsu). These polymerases both lack 3′-5′ exonuclease activity. LF-Bsu has been modified to remove the exonuclease activity that Pol IV intrinsically lacks. In the following, we will present experimental results for both proteins indicating that under conditions relevant *in vivo*, DNA synthesis initiated by RecA-mediated homology recognition is highly unlikely unless there is a sequence matched heteroduplex product with length L_prod_ > 50 bp that terminates within 8 bp of the 3′ end of the initiating strand.We first formed ssDNA-RecA filaments and then allowed these filaments to interact with the dsDNA. If a sufficiently stable heteroduplex forms, a DNA polymerase can extend the initiating strand.

We first 150 formed ssDNA-RecA filaments and then allowed these filaments to interact with the dsDNA. If a sufficiently stable heteroduplex forms, a DNA polymerase can extend the initiating strand. That extension begins at the terminal 3′ OH of the initiating strand and proceeds in the 5′ to 3′ direction with respect to the initiating strand. In our *in vitro* experiments, the synthesis can eventually reach an end of the dsDNA. We will refer to that end of the dsDNA as the 3p end. We will specify positions in the dsDNA using D, their separation from the 3p end of the dsDNA. We monitored the base pairing between the two strands in the dsDNA by measuring the emission due to a fluorescein label on one of the dsDNA strands (Figure 3a). Initially, the fluorescein emission is quenched by the nearby rhodamine label on the other strand, but if dsDNA separates, the fluorescence emission will increase.

**Figure 3.**
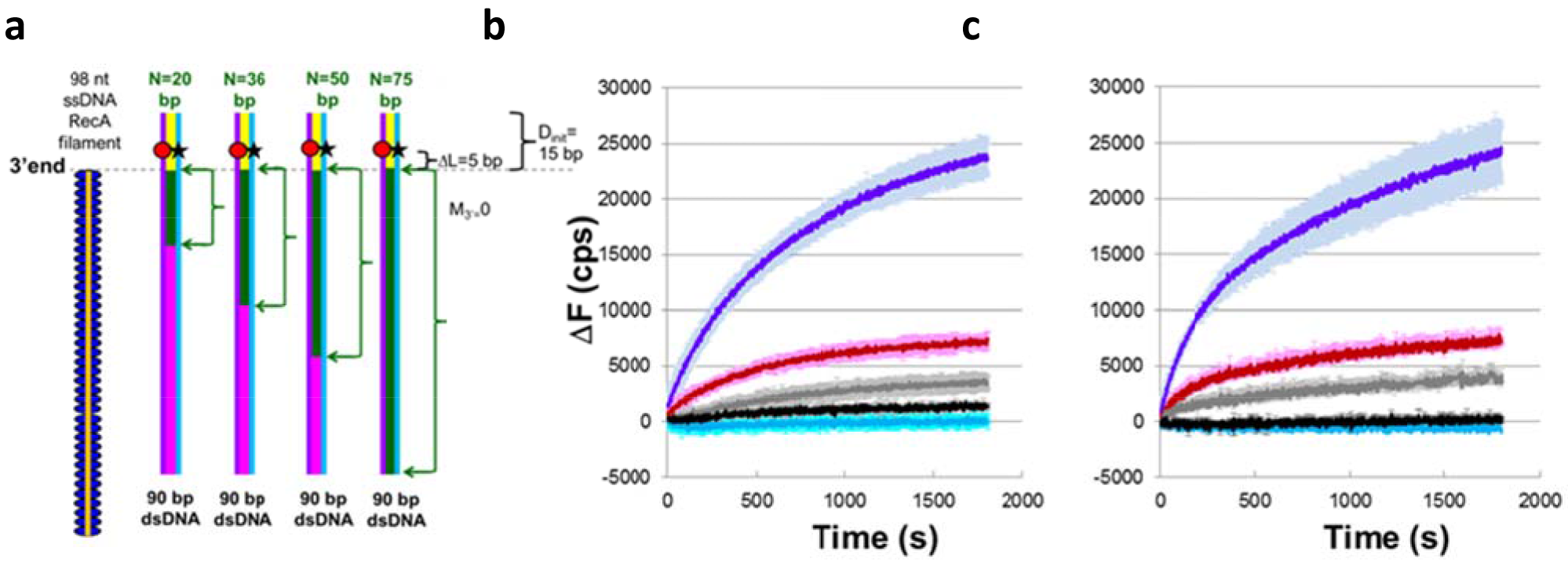
DNA synthesis stabilizes DSB repair occurring at a repeat. **a** Experimental schematic showing a typical ssDNA-RecA filament (orange line with blue ellipses) and dsDNA. ΔL= D_label_-D_init_ = 5 bp. The labeled dsDNA used in all of the experiments was the same, so each N value corresponds to a different filament sequence. For each N value, the green arrows highlight the green regions of the dsDNA that are homologous to the N bases at the 3′ end of the ssDNA. The other bases in the dsDNA are heterologous to the initiating ssDNA. The yellow region indicates D_init_ = 15 bp. The remaining dsDNA is shown in magenta. The red circle and black star represent the rhodamine and fluorescein labels, respectively. They are positioned on the complementary (purple line) and outgoing (blue line) strands, respectively. **b** Graph representing the average over three trials of the change in fluorescence (AF) vs. time curves in experiments with dATP-ssDNA-RecA filaments and DNA Pol IV represented in (**a**) for N = 75 (dark blue), 50 (red), 36 (gray), 20 (black), and heterologous filament (light blue). AF in counts per second (cps) is calculated as the difference between the measured fluorescence and the average initial fluorescence for heterologous dsDNA. The error bars show the standard deviation based on three trials. **c** Same as (**b**) in the presence of ATP-ssDNA-RecA filaments and LF-Bsu polymerase.

To study effects due to the DNA polymerases, we positioned the dsDNA labels ΔL base pairs beyond the 3′ end of the filament. ΔL was chosen to be large enough that long strand exchange products do not produce significant fluorescence increases even if the product extends to the 3′ end of the filament. In what follows, we will show that under these conditions the presence of a DNA polymerase lacking 3′-5′ exonuclease activity can produce large fluorescence increases as long as RecA filaments and dNTPs are present. Importantly, this fluorescence also depends strongly on N, the number of contiguous bases in the dsDNA that are complementary to the corresponding bases in the initiating strand.

In the first set of experiments, the fluorescent labels were located at D_label_ ~10 bp and the 3′ end of the initiating strand was positioned at D_init_ = 15 bp as shown schematically in Figure 3a (bracket on top of the schematic). The 15 base pairs that extend beyond the 3′ end of the filament are indicated in yellow. The same 90 bp labeled dsDNA target was used in all of the experiments illustrated in Figure 3a. We varied the homology between the dsDNA and the ssDNA-RecA filaments by changing the sequence of the initiating ssDNA. In particular, different 98 nt sequences were designed to be heterologous to the dsDNA except for N contiguous bases at the 3′ end of the filament that match the corresponding N bases in the dsDNA (shown by the green brackets in Figure 3a, and encompassing 20, 36, 50, and 75 base pairs).

Figure 3b shows graphs of ΔF, the difference between the measured fluorescence as a function of time and the average initial fluorescence value for a heterologous ssDNA-RecA filament. These experiments were carried out with DNA Pol IV, ssDNA-RecA filaments, and dNTPs. Figure 3c shows the analogous results with LF-Bsu. In Figure 3b, c, each of the curves represents results for different N values. Results obtained without DNA polymerase are shown in Figure 3- figure supplement 1, along with results obtained with DNA Pol IV and RecA, but without dNTPs. Comparison of Figure 3b, c with Figure 3- figure supplement 1 suggests that the observed fluorescence increase is dominated by DNA synthesis rather than dsDNA melting due to either strand exchange alone or DNA Pol IV binding without synthesis. Thus, those results suggest that in experiments performed with dNTPs, the fluorescence signals are dominated by DNA synthesis that extends the initiating strand toward the fluorescent labels.

If the DNA synthesis that dominates the contribution to that fluorescent signal made recombination irreversible, the curves for N = 75 and N = 50 in Figure 3 would approach the same asymptotic value corresponding to 100 % product formation. In contrast, Figure 3b, c shows that the results for N = 50 approach a lower asymptotic value than the results for N = 75. The significant but lower asymptotic value achieved by N = 50 suggests that synthesis that is sufficient to trigger observable fluorescence does not always create a product in which the initiating and complementary strands are irreversibly paired. Thus, Figure 3 shows that the complementary strand can return its base pairing to the outgoing strand even after some synthesis has occurred.

Additional results for LF-Bsu are shown in Figure 3- figure supplement 2. In those experiments Dinit is also 15 bp, but the fluorescent labels are positioned at the 3′ end of the filament (D_label_=0 and ΔL = 15), whereas in Figure 3 D_label_ = 10 and ΔL = 5. Those results also indicate that the fluorescence increase due to DNA synthesis is small unless N > ~36 bp. In sum, even though the intrinsic processivity for DNA Pol IV is different from the processivity of LF-Bsu, the similarity between Figure 3b, c and Figure 3- figure supplement 2 suggests that the results represent general features of DNA synthesis triggered by the formation of heteroduplex products, at least for DNA polymerases that lack 3′ to 5′ exonuclease activity.

### Adjacent homoduplex dsDNA decreases product stability

*In vivo,* the three strand heteroduplex products resulting from the pairing of the initiating and complementary strands are almost always flanked by homoduplex dsDNA of the complementary and outgoing strands. Previous work has suggested that this homoduplex dsDNA drives reversal of adjacent heteroduplex products (Danilowicz et al., 2017). To probe the importance of these molecular events, we increased D_init_ from 15 bp to 66 bp because D_init_ is equal to the number of bases that must be synthesized to traverse the homoduplex dsDNA so it becomes fully separated at the 3′ end of the initiating strand. If strand displacement synthesis by a DNA polymerase is rapid enough to reach the 3p end of the dDNA when D_init_ = 15, but not rapid enough to reach the end when D_init_ = 66, then comparison of results from experiments with the two different D_init_ values may provide insight into the influence of homoduplex dsDNA adjacent to the three strand heteroduplex products.

The same dsDNA with D_label_ = 58 bp was used in all of the experiments illustrated in Figure 4a, and N was controlled by varying the 98-nt sequence of the initiating strands. For this construct, even for N = 82, we see no increase in fluorescence in the absence of DNA synthesis. The raw fluorescence curves obtained with dATP-ssDNA-RecA filaments, DNA Pol IV, and dNTPs are shown in Figure 4- figure supplement 1, and Figure 4b shows the graphic representation of the corresponding change in fluorescence, ΔΔF vs time curves, where ΔΔF is the difference between the observed fluorescence and the fluorescence for N = 5 at each time. In the figure, the purple, red, and black curves represent results for N = 82, 50, and 20, respectively. As shown in Figure 4- figure supplement 1, the increase in fluorescence is only statistically significant if N > ~50. Figure 4- figure supplement 2 shows analogous results for ATP-ssDNA-RecA filaments, LF-Bsu, and dNTPs. Comparison of Figure 3b (D_init_ = 15, AL = 5), Figure 3- figure supplement 2 (D_init_ = 15, AL = 15), and Figure 4b (D_init_ = 66, AL = 8), indicates that adjacent homoduplex regions destabilize heteroduplex products even in systems that include DNA synthesis.

**Figure 4.**
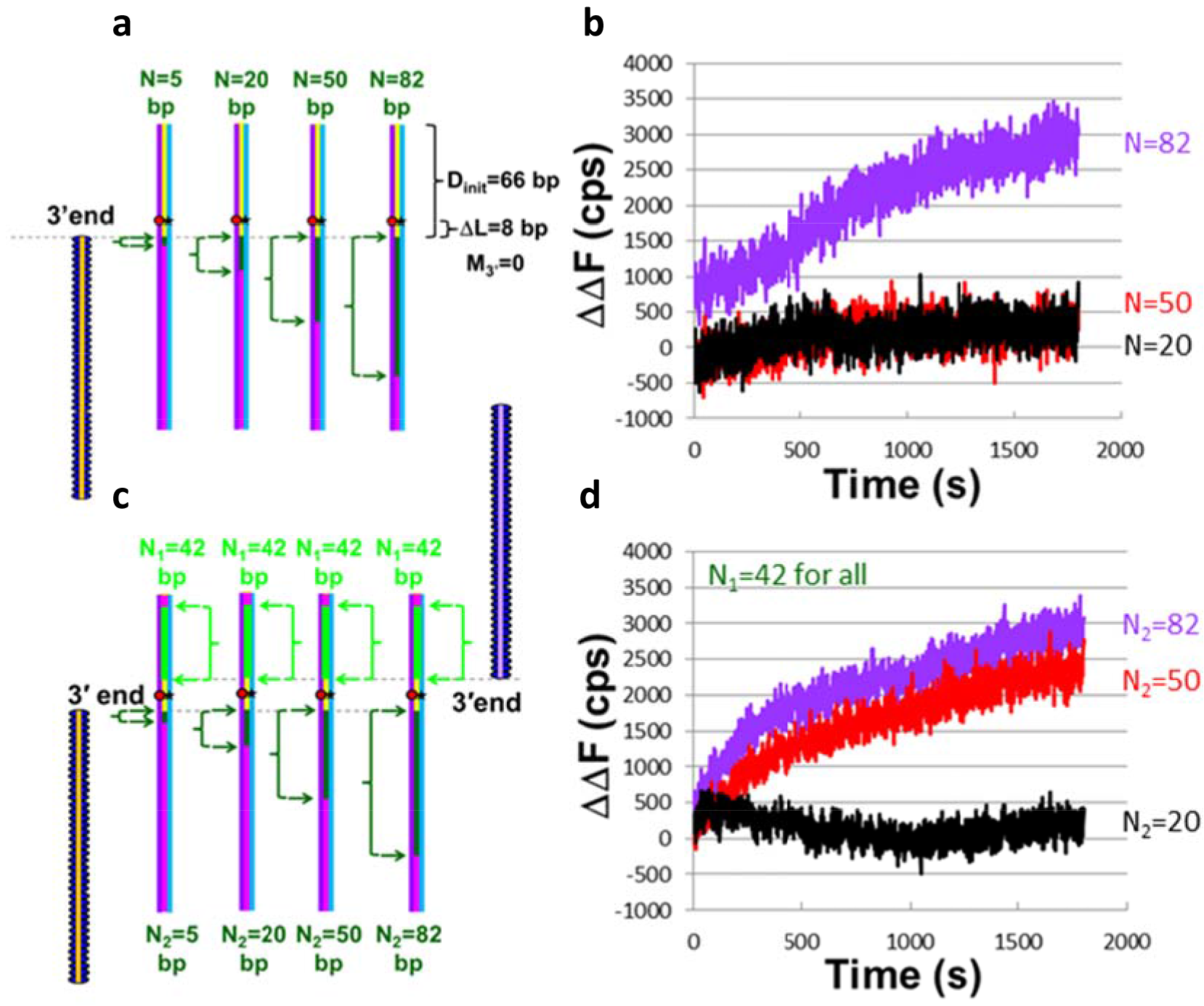
The presence of a second filament rescues instability caused by an ssDNA outgoing strand and a long homoduplex dsDNA that extends beyond the 3′ end of a filament. **a** Schematic of experiments with 66 bp of homoduplex dsDNA on the 3′ end of the filament and N values of 5, 20, 50, and 82 bp. b Graphic representation of the change in fluorescence (ΔΔF) vs. time curves of the experiment represented in (**a**) with dATP-ssDNA-RecA filaments and DNA Pol IV. ΔΔF is calculated as the difference between the measured fluorescence and the fluorescence for N = 5. The black, red, and purple curves correspond to N = 20, 50, and 82, respectively. **c** Schematic for experiments performed involving two filaments. In all of these experiments N_1_ = 42. **d** Graphic representation of the change in fluorescence AA1F vs. time curves for single trials of the experiment represented in c with dATP-ssDNA-RecA filaments and DNA Pol IV. ΔΔF was calculated as indicated above. Different N_2_ values are represented with different curve colors, where purple, red, and black correspond to N_2_ = 82, 50, and 20 nt, respectively.

### Synthesis triggered by two filaments stabilizes recombination products

As illustrated in Figure 1, if each of the filaments triggers DNA synthesis that completes a double strand, then no unpaired bases will remain. To study synthesis triggered by the initiating ssDNA formed at both sides of a DSB, we performed the experiments illustrated in Figure 4c. All of the experiments illustrated in Figure 4c included one filament with N_1_ = 42 contiguous bases that are homologous to the corresponding bases in the dsDNA. The sequence of the second filament was varied so that N_2_, the number of contiguous bases that are homologous to the corresponding bases in the other strand of the dsDNA, varied from 0 to 82 bases. The ΔΔF results shown in Figure 4d, analogous to results shown in Figure 4b, indicate that the fluorescence change for N_2_ = 50 is quite significant, even though no detectable fluorescence change was observed in one-filament experiments with N = 50 (Figure 4b); therefore, a second filament with N_1_ = 42 significantly increased the fluorescence shift observed for filaments with 50 contiguous homologous bp. Thus, comparison of Figure 4b and 4d indicates that a second initiating ssDNA significantly increases the probability that the outgoing and complementary strands will be separated in the region between the filaments. This must be the result of a cooperative interaction between the two filaments because the signal due to either individually was negligible.

This cooperative increase in product stability is consistent with both filaments triggering synthesis within the dsDNA region containing the labels, resulting in both the fluorescein and rhodamine labels being incorporated in different dsDNA strands. As a result, the restoration of quenching is less likely than it would be if one of the labeled strands remains unpaired and available to pair again with its original partner (Rosselli and Stasiak, 1990, Danilowicz et al., 2017). Figure 4- figure supplement 2 shows that when synthesis is performed by LF-Bsu, the presence of a second filament with NΔΔ = 42 significantly enhances ΔΔF, if N_2_ is at least 20 bp even though Pol IV required N_2_ = 50 bp. The difference in the required N_2_ values for the two polymerases may reflect the more efficient strand displacement synthesis provided by LF-Bsu. Furthermore, even for the case where N_2_ = 82 bp, the results for DNA polymerase Pol IV that are shown in Figure 4d are ~ 4× smaller than the fluorescent values for LF-Bsu that are shown in Figure 4- figure supplement 2. The much smaller fluorescent values obtained for DNA polymerase Pol IV suggest that product formation in this case is low.

As shown in Figure 2e and 2f, if the system followed the RecBCD pathway, the separation between the 3′ ends of the filaments will be much longer than the 16 bp separation used in these experiments. Thus, *in vivo,* formation of irreversible products triggered by initiating strands with ~ 40 bp N_1_ and N_2_ is probably much smaller than the low product formation shown in Figure 4d because *in vivo* the separation between the 3′ ends of the two filaments is so much longer than 16 bp; however, comparison of Figure 4b, d does show that product stability can increase greatly if both initiating strands trigger synthesis that creates regions in which all of the DNA strands are base paired.

For the experiments shown in Figures 3b, c, 4b, d, Figure 3- figure supplement 2, and Figure 4- figure supplement 1 and 2, the N contiguous homologous bases are positioned at the 3′ end of the filament, but we also wanted to explore cases in which M_3′_ mismatches separated the N sequence matched bases from the 3′ end of the filament. We performed M_3′_ > 0 experiments to determine whether pairings between long repeats that are distant from the 3′ end of the filament could trigger genomic rearrangement since *in vivo* heterologous dsDNA always surrounds sequence matched heteroduplex products formed by joining different copies of long repeats.

Like eukaryotic recombinases, RecA can create strand exchange products that include some mismatches (Volodin et al., 2009, Sagi et al., 2006). However, there is also evidence indicating that the efficiency of strand exchange decreases in the presence of mismatches (Danilowicz et al., 2015, Qi et al., 2015). Thus, extension of the heteroduplex to the 3′ end of the filament may become increasingly improbable as the number of mismatches at the 3′ end of the filament increases. Since DNA synthesis triggered by strand exchange extends the initiating strand using the complementary strand as a template (Pomerantz et al., 2013), the synthesis requires the DNA polymerase to interact with the heteroduplex and the 3′ OH at the end of the initiating strand. Thus, DNA synthesis is likely improbable if the heteroduplex product rarely incorporates the mismatched bases at the 3′ end of the filaments.

### Synthesis is blocked by mismatches at the 3′ ends of ssDNA

To test whether mismatches at the 3′ end of the filament can inhibit the DNA synthesis required to make recombination irreversible, we designed experiments to study how M_3′_, the number of mismatches at the 3′ end of the filament, influences the interaction between the strand exchange product and the DNA polymerase. The experiments are illustrated schematically in Figure 5a. Figure 5b shows the ΔF curve obtained in the presence of DNA Pol IV and indicates that even M_3′_ = 3 strongly suppresses the fluorescence increase, suggesting no strand separation due to DNA synthesis. Furthermore, the result for M_3′_ = 5 is indistinguishable from the results for heterologous controls. Analogous results obtained in the presence of LF-Bsu show that even M_3′_ = 3 (Figure 5c) is indistinguishable from the heterologous controls (Figure 5- figure supplement 1). Additional experiments were performed with N = 82 and the construct illustrated in Figure 4a. Figure 5- figure supplement 2 shows results for experiments with N = 82 and either M_3′_ = 0 or M_3′_ = 8. Controls with M_3′_ = 0 and either N = 5 or N = 0 are also shown; results for N = 82 and M_3′_ = 8 are indistinguishable from the heterologous controls. For that system, lower M_3′_ values were not tested.

**Figure 5.**
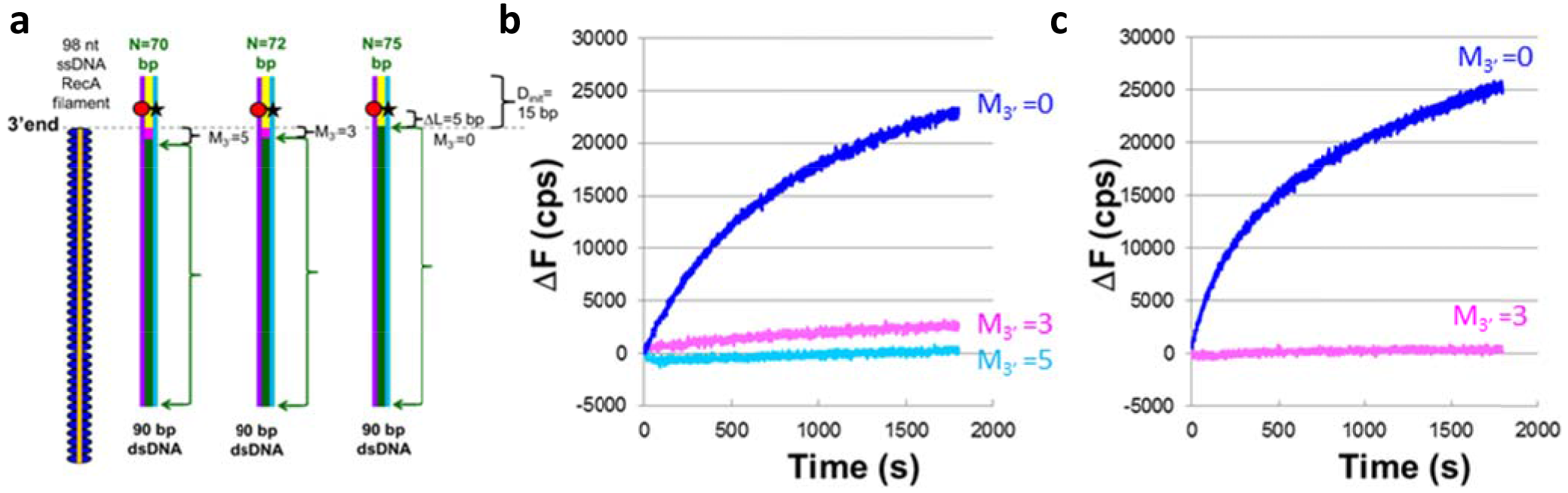
DNA synthesis is required for a DNA Pol to stabilize strand exchange products. **a** Schematic for experiments with M_3′_ values of 0, 3, 5. **b** Graphic representation of the change in fluorescence (ΔF) vs. time curves from single trial experiments performed with dATP-ssDNA-RecA filaments and DNA Pol IV in which the blue, pink, and light-blue curves correspond to M3′ values of 0, 3, and 5 base mismatches, respectively. ΔF is calculated as the difference between the measured fluorescence and the average initial fluorescence for heterologous dsDNA. **c** Analogous experiments as (**b**) but with LF-Bsu polymerase instead of DNA Pol IV.

### Chi sites rarely occupy the 3′ ends of long repeats

We will refer to the sequence provided by the genome database as the “given” strand. The other strand in the genome is complementary to the given strand, so we refer to that strand as the “comp” strand. In the RecBCD pathway, as indicated in Figure 1aiv, one initiating ssDNA will terminate with a Chi site from the given strand and the other initiating ssDNA will terminate with a Chi site from the comp strand.

Figure 6 displays graphical information designed to highlight the positions of Chi sites in long repeats and the decrease in repeat length due to RecBCD. In particular, Figure 6 a, b shows all of the repeated sequences in *E. coli* O157 that are longer than 20 bp and include at least one Chi site. The total height of the bars in Figure 6 a, b represents N_repeat_, the total length of the repeat. No repeat includes a Chi site on both strands, so we have separated the results according to the strand in which the Chi sites appear. The upper end of each bar corresponds to the 3′ end of the strand. An expanded view of these figures is shown in Figure 6- figure supplement 1.

**Figure 6.**
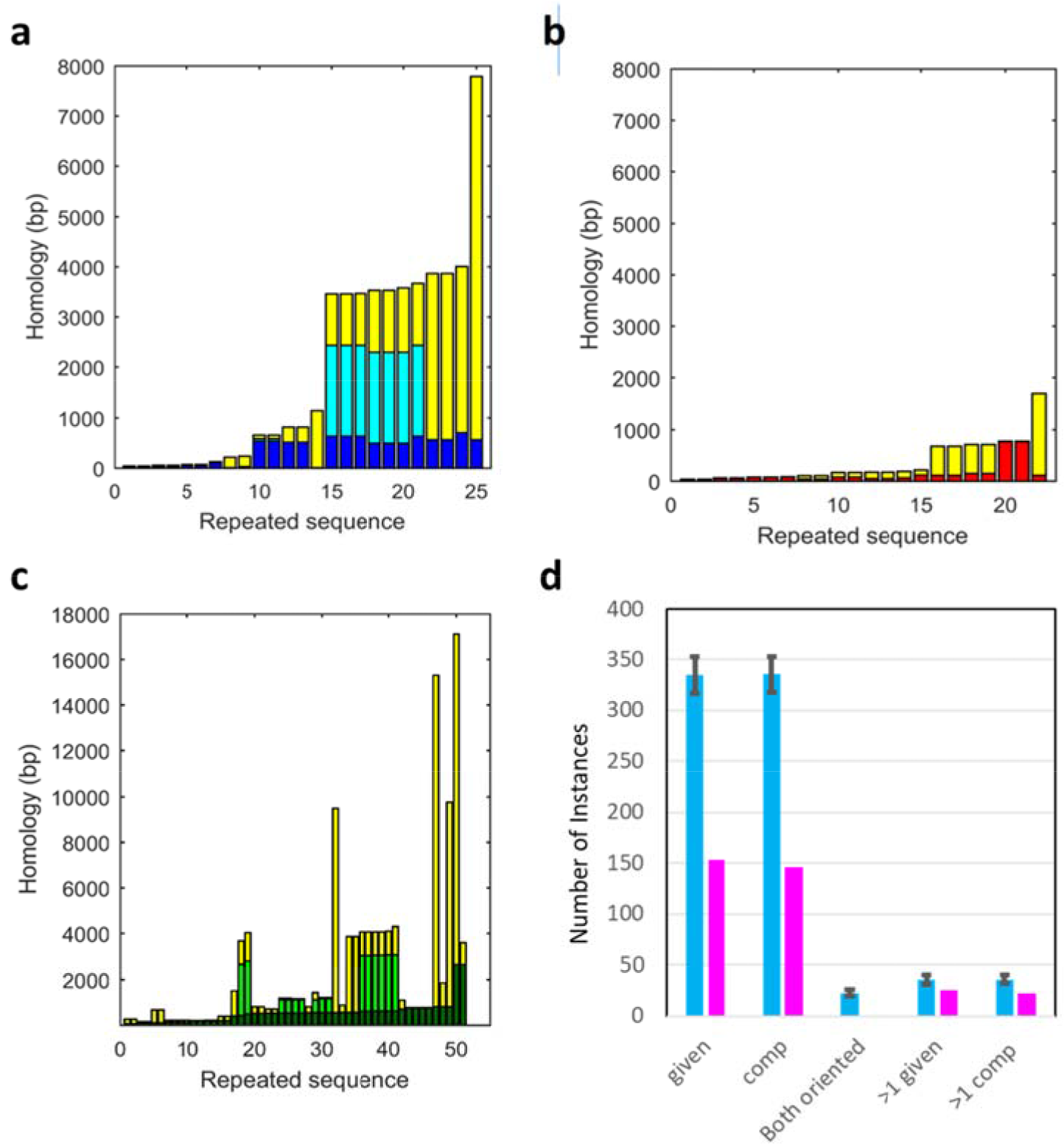
Chi sites in *E. coli* O157 are rarely located at the 3′ end of long repeated sequences. **a** Same as Figure 2a but showing only repeats in the given strand that include a Chi site. The total height of the bar corresponds to the total length of the repeat. The height of the yellow regions corresponds to the region that would be always degraded by RecBCD. The height of dark blue regions corresponds to the separation between the 5’ end of the repeat and the 3’ end of the nearest Chi site in the repeat. For repeats that contain two Chi sites, the region between the Chi sites is shown in cyan. No repeat includes more than two Chi sites. **b** Same as (**a**); analogous data for the comp strand with regions on the 5′ side of the Chi site shown in red. **c** Filaments including repeats > 60 bp summed over both strands in 4 *E. coli* genomes. The dark green regions show the separation between the 5′ end of the repeat and the nearest Chi site. The light green regions indicate the spacing between Chi sites. **d** The height of the magenta bars indicates the number of Chi sites in repeats > 20 bp, and the blue bars represent the corresponding results for markers randomly positioned in the genome, where the number of markers/strand is equal to the number of Chi sites/strand. The results are summed over 12 enteric bacteria. The first two sets of bars show results for repeats in the given strand and the comp strand, respectively. The next set of bars show cases in which one repeat would create two filaments, each of which include > 20 bp from the repeat. For Chi sites, this never occurs. The final two sets of bars show the number of cases in which more than one marker or Chi site is positioned in the same repeat on the same strand. The error bars represent the standard deviations for the randomly positioned markers.

Each bar is divided into colored regions that represent the relationship between that region and the 3′ end of Chi sites. The yellow regions indicate portions of the repeats that are on the 3′ side of all of the Chi sites in the repeat, so the yellow regions do not participate in any homology search. The separation between the 5′ end of the repeat and the 3′ end of the nearest Chi site is shown in dark blue and red, for the given and comp strands, respectively. These regions would participate in the homology search if RecBCD recognizes the Chi site nearest the 5′ end. Though no comp strand repeat in this genome contains more than one Chi site, seven repeats in the given strand contain two Chi sites. The cyan bar shows the separation between the two Chi sites.

Figure 6c shows analogous results averaged over both strands in four *E. coli* genomes, restricted to cases where the 5′ end of the repeat is separated from the 3′ end of the nearest Chi by > 60 bp. In Figure 6c, the green regions indicate the number of bp on the 3′ side of all Chi sites, and the dark green regions are analogous to the red and dark blue regions in Figure 6 a, b. Light green indicates regions between two Chi sites in the same repeat. No repeat contained more than two Chi sites. The figure shows that for repeats with lengths <~1000 bp, the positioning of the Chi sites within the repeats allows RecBCD to reduce the length of the repeat that participates in the homology search, which *in vivo* data suggests reduces rearrangements due to joining different copies of the repeat (Bi and Liu, 1994).

If Chi sites play a role in avoiding recombination due to interactions between long repeats, then one would expect that the number of Chi sites positioned in long repeats would be suppressed with respect to a system in which the Chi sites were randomly positioned in the genome. To test this, we randomly positioned markers within each strand of real genomes, where the number of markers in each strand was equal to the number of Chi sites in that strand. The results shown in Figure 6d indicate that Chi sites positioning in long repeats is strongly suppressed. The detailed data for each genome are shown in Supplementary Table 1.

In calculating the results in Figure 6, we only considered repeats that included at least 20 bp on the 5′ side of the Chi site, which would be the interaction if all of the bases in the Chi site were degraded. We note that the results shown in Figure 6 are not significantly altered if Chi site occupies the 3′ end of the filament since allowing the Chi site to remain only adds one new repeat pair to the comp strand and adds one additional occurrence to two 20 bp sequences that were already repeated twice.

### The fraction of DSB creating filaments ending in long repeats

As shown in Figure 1b, in the hypothetical DSB repair mechanism, the sequences at the 3′ ends of the filaments flank the DSB, so the filament sequences uniquely specify the position of the DSB that created the filaments. In contrast, as shown in Figure 1a, if the RecBCD pathway is followed, the same sequences at the 3′ ends of the filaments can result from any DSB positioned between Chi sites on the 3′ ends of the two filaments (i.e. L_chi_). Thus, the effectiveness of the RecBCD pathway in reducing genomic recombination cannot be determined by simply considering how many Chi sites have repeated sequences at the 5′ side.

Additional information can be gained by considering all of the possible DSB positions in the genome and determine what fraction of them lead to initiating ssDNA with long repeated sequences at 9their 3′ ends. Importantly, no long repeated sequence that appears on the 5′ side of a Chi site appears elsewhere in the genome without the adjacent 8 bp Chi site. Thus, if even all of the Chi site bases are degraded before the searching filament is formed, in the RecBCD pathway genomic rearrangement can only occur by joining long repeats that occupy the 5′ side of a Chi site. For these calculations, we assumed that DSBs are distributed randomly on the genome and that the function of RecBCD is changed by the first Chi site it encounters. Given these assumptions, we calculated the fraction of the DSBs that create initiating strands whose 3′ ends terminate in at least one repeat containing N_rep 3′_ > n bases on a specified initiating strand (DSB1_frac_(n)) or on both initiating strands (DSB2_frac_(n)).

Figure 7a shows the results for the RecBCD pathway. To calculate the results, we first computed DSB1_frac_(n) for each strand in each of 12 enteric bacteria that have the same Chi site sequence (5′-GCTGGTGG-3′)^41^.We then averaged the results for each strand over all of the 12 bacteria to get the average probabilities for each strand. The red and blue lines in Figure 7a show DSB1_frac_(n) for the given and comp strands, respectively. They represent the probability that a DSB will lead to the formation of a filament from each strand with N_rep 3′_ exceeding the x-axis value. The black line shows the sum of the two probabilities. The graph indicates that ~ 2 % of all DSB would create at least one filament with a repeat on its 3′ end that could pass a 300 bp homology test. This suggests that substantial genomic rearrangement could occur if only one filament was required to pass the homology test; however, strand exchange products remain reversible (Rosselli and Stasiak, 1990, Danilowicz et al., 2017) unless two complete dsDNA strands are formed. Formation of two complete dsDNA requires that both searching filaments trigger synthesis. If a DSB forms within a repeat, major genomic rearrangement will result if both searching filaments pair with another copy of that repeat.

**Figure 7.**
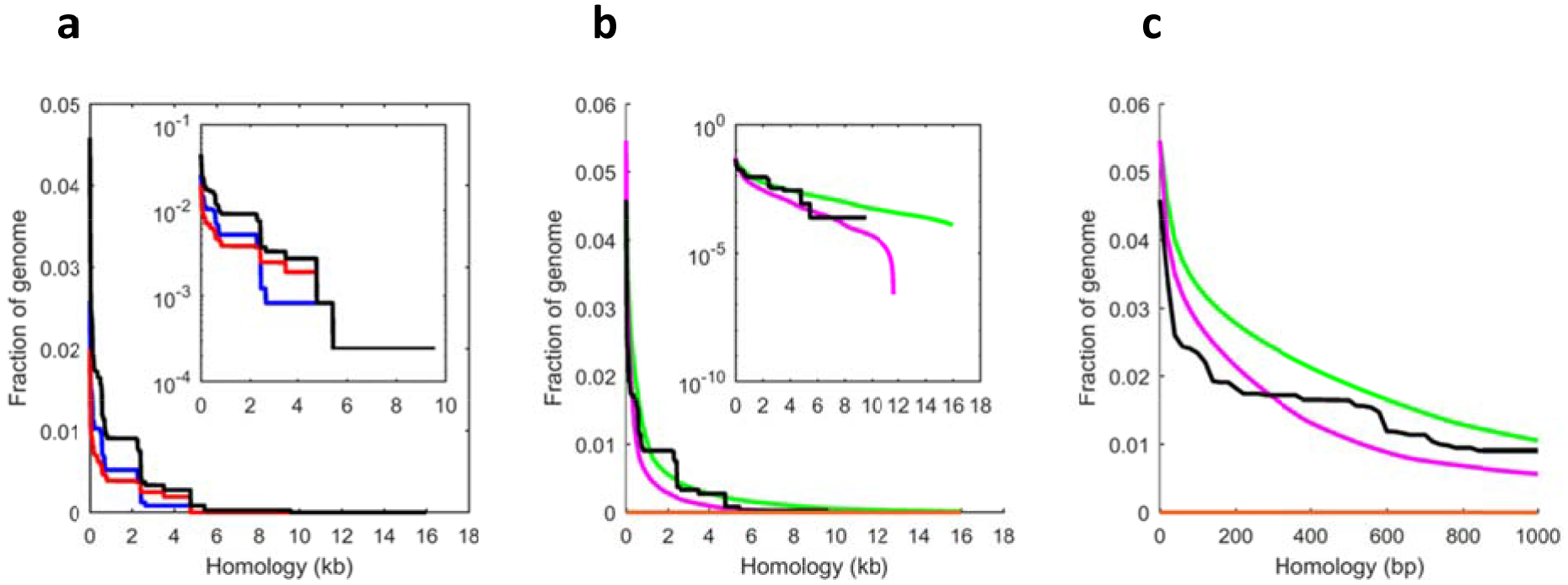
Filaments created by the RecBCD pathway do not end in long repeated sequences. **a** All possible DSB1(_frac_)(n) and DSB2(_frac_)(n) during RecBCD-mediated DSB repair of twelve enteric bacterial genomes. The blue and red curves show DSB1_frac_(n) for the given and comp strands, respectively. The black curve shows the sum of the red and blue curves. The horizontal orange line along the x axis shows DSB2_frac_(n) = 0 for all n. For all curves the bin sizes are 20 bp. The inset shows the same data with an expanded x axis and a logarithmic y axis. **b** The black and orange lines are the same as in (**a**). The green and magenta curves show the analogous result for the hypothetical DSB ends mechanism. The inset shows the same data with an expanded x axis and a logarithmic y axis. **c** Same as (**b**) but with an extended x axis.

Thus, we considered DSB2_frac_(n), and found that no genome contained a repeat that could create N_rep 3′_ > 20 bases on both initiating strands, as indicated by the orange line that lies along the x-axis in Figure 7a-c. For *E. coli* O157, we also considered the 8 cases in which different copies of a repeat contained Chi sites on opposite strands; however, in all cases those two Chi sites were separated by more than 15 Chi sites on either strand, so given the 30 % probability of recognizing a Chi site, it is enormously unlikely one DSB would produce two filaments terminating in those Chi sites.

Figure 7b, c highlights some advantages of the RecBCD pathway by comparing the results shown in Figure 7a to the results for the hypothetical DSB ends mechanism. The black and orange lines in Figure 7b, c are the same as those in Figure 7a, but Figure 7b, c also shows green and magenta curves representing the analogous results for the hypothetical DSB ends mechanism. The difference between the green and black curves provides some information about the influence of Chi sites on genomic rearrangement as a result of suppression of DSB1_frac_(n), but rearrangement probabilities are also influenced by the number of times a repeat occurs in the genome and the physical distance between the repeat at the end of the filament and other copies of the repeat, where that distance may change with time.

Fortunately, the difference between the magenta and orange is much easier to interpret because the orange line indicates that if the RecBCD pathway is followed no DSB would create two filaments that would include regions of the same repeat. In contrast, in the DSB ends pathway many do. Importantly, Figure 6d shows that summing over the results for all 12 genomes yielded > 20 instances in which two Chi sites on the same strand occur in one repeat. That statistic predicts that summing over the same genomes should yield ~ 20 repeats that could create N_rep 3′_ > 20 bases on both initiating strands; however, the actual sum was zero; consequently, for the RecBCD pathway the suppression of DSB2_frac_(n) is not the result of the observed reduction of instances in which Chi sites occupy one strand on a repeat. Thus, the statistical distribution of Chi sites in the genomes of enteric bacteria suggests that strong suppression of DSB2_frac_ is much more important than preventing Chi sites from occupying one strand in a repeat. This strong suppression avoids formation of searching filament pairs that include regions of the same long repeat at their 3′ ends, so the strong statistical suppression supports our proposal that the placement of Chi sites allows the RecBCD pathway illustrated in Figure 1a to strongly suppress genomic rearrangement; however, it is probable that in rare instances Chi sites may be associated with increased genomic recombination if the system does not follow the pathway shown in Figure 1a.

## Discussion

In sum, it has been known for decades that homologous recombination in bacteria frequently occurs at Chi sites, which are significantly overrepresented (~10x random probability) in bacterial genomes; however, the benefits conferred by Chi sites had remained elusive. In this work, we have presented experimental and theoretical evidence for an elegant mechanism that exploits the sequence distributions near Chi sites to suppress genomic rearrangements that would otherwise be both frequent and fatal. We note that eukaryotic genomes are longer and contain more long repeated sequences, so a “Chi site” system that includes only 8 bases might not be effective in longer genomes. Of course, it is possible that eukaryotes could use a similar system involving more than 8 bases, or 8 base sequences might provide some rejection if double strand break repair in eukaryotes was confined to domains that included only a few Mbp.

## Methods and Materials

### FRET measurements

Strand exchange reactions were performed by mixing an aliquot of 0.06 μM 98 nt ssDNA/RecA filament, 0.06 μM labeled dsDNA, and 1 μM *E. coli* DNA Polymerase IV (obtained using DinB overproducer plasmids (Tashjian et al., 2017, Cafarelli et al., 2013) or 5 units Bsu DNA polymerase, Large fragment (LF-Bsu) (New England Biolabs (NEB), 5000 units/ml) and rapidly transferring the solution to a quartz cuvette. For DNA Pol IV measurements, the RecA buffer contained 0.1 mg/ml BSA, 2 mM dATP, and 0.4 mM dNTPs. Measurements in the presence of Bsu polymerase were performed in RecA buffer containing 1 mM ATP and 0.1 mM dNTPs.

The filaments were initially prepared by incubating 0.06 μM ssDNA (final concentration ~6 μM in bases) with 2 μM RecA (NEB) in the presence of 1 mM cofactor (ATP or dATP), 10 U/ml of pyruvate kinase, 3 mM phosphoenolpyruvate, and 0.2 μM single-stranded binding protein (SSB) in RecA buffer (70 mM Tris-HCl, 10 mM MgCl_2_, and 5 mM dithiothreitol, pH 7.6) at 37°C for 10 minutes.

FRET experiments followed the emission of the fluorescein label by using 493-nm excitation during 30 minutes; the emission was read as counts per second (cps) at 518 nm every one second. The integration was 0.5 s and the band width 2 nm. The sample was kept at all times at 37°C.

The dsDNA containing 90 bp with internal labels was obtained by heating and cooling down slowly the corresponding oligonucleotides from 90 to 40°C with 1°C steps equilibrated for 1 minute; the emission at 518 nm was acquired (excitation at 493 nm) at each temperature step.

The dsDNA containing 180 bp was prepared by initially annealing a 90 nt ssDNA containing an internal rhodamine label on base 58 from the 5′ end and a 5′-end phosphorylated oligonucleotide (82 bases) containing an internal fluorescein label (position 57 from the 3′ end). Another dsDNA without labels was annealed using two oligonucleotides containing 90 and 98 bases; the former was 5′-end phosphorylated. Finally the two dsDNAs were annealed and ligated overnight at 16°C in the presence of T4 DNA ligase in ligase reaction buffer (50 mM Tris, 10 mM MgCl_2_, 1 mM ATP, and 10 mM dithiothreitol, pH 7.5, NEB). The 180 bp construct was further purified by running a 3 % agarose gel in TBE (Tris/Borate/EDTA) buffer for 2 hours (6 V/cm). The 180 bp band was visualized with a midrange UV trans-illuminator and cut. Finally the dsDNA was extracted from the agarose using a Nucleospin kit (Machery and Nagel, Bethlehem, PA) and concentrated on a YM-100 centrifugal filter (Millipore).

The sample containing 98 bp dsDNA was prepared by annealing the complementary oligonucleotides from 90 to 40°C with 1°C steps equilibrated for 1 minute; the emission at 518 nm was acquired (excitation at 493 nm) at each temperature step.

### Oligonucleotides used for dsDNA preparations and filaments

Oligonucleotides for dsDNA 90 bp with internal fluorophores: 5′ CGG AAA TCA C/iRho-T/C CCG GGT ATA TGA AAG AGA CGA CCA CTG CCA GGG ACG AAA GTG CAA TGC GGC ATA CCT CAG TGG CGT GGA GTG CAG GTA 3′ and 5′ TAC CTG CAC TCC ACG CCA CTG AGG TAT GCC GCA TTG CAC TTT CGT CCC TGG CAG TGG TCG TCT CTT TCA TAT ACC CGG GAG /iFluor-T/GA TTT CCG 3′.

Oligonucleotides for filaments interacting with 90 bp dsDNA. 75 (−15) plus 23 heterologous: 5′ GGACA CTGCTTCATTCCTCTTATTACCTGCACTCCACG CCACTGAGGTATGCCGCATTG CACTTTC GTCCCTGGCAGTGGTCG TCTCTTTCATATACC 3′; 50 (−15) plus 48 heterologous: 5′ GGACGCT GCCGGAT TCCTGTTGAGTTTATTGCT GCCGTCATTGCTTATATGCCGCAT TGCAC TTTCGT CCCTGGCAGTGGTCGTCTCTTTCATATACC 3′; 36 (−15) plus 62 heterologous: 5′ GGACGCTGCC GGATTCCTG TTGAGTTTATTGCTGCCGTC ATTGCTTATTATGTTCA TCCCG TTTTCGTCCC TGGCAGTGGTCGTCTCTTTCATATACC 3′; 20 (−15) plus 78 heterologous: 5′ GGACGCT GCCGG ATTCCTGTTGAGTTTATTGCTGCCGTCATTGCTTATTATGTTCATCCCGTCAACATTCAA ACGGCCGGTCGTCTCTTTCATATACC 3′; 3 mismatches 3′ end: 5′ GGACGCTGCCGGATTCCTG AGTATACCTGCACTCCACGCCACTGAGGTATGC CGCATTGC ACTTTCGTCCCTGGCAGT GGTCGTCTCTTTCATATTAA-3′; 5 mismatches 3′ end: 5′ GGACGCTGCCGGATTCCTCTGTATA CCTGCACTCCACGCCACTGAGGTATGCCGCATTGCACTTTCG TCCCTGGCAGTGGTCGTCT CTTTCATTCTAA 3′

### Oligonucleotides for 180 bp dsDNA

Annealed initially with labels: 82 nt (Flu57): 5′(K) CTCCACGCCACTGAGGTATGCCGCA/iFluorT/ TGCACTTTCGTCCCTGGCAGTGGTCGTC TCTTTC ATATACCCGGGAGTGATTTCCG 3′ and 90 nt (Rho58): 5′ CGGAAATCACTCCCGG GTATATGA AAGAGACGACCACTGCCAGGGACGA AAGTGCAA/iRhoT/GCGGCATACCTCAG TGGGTGGAGTGCAGGTA 3′. Annealed initially (no labels): 90 nt: 5′ (K)AATCCGGCAGCGTCCGTCGTTGTTGATATTGCTTATGAAGGCTCC GGCAGTGGCGACTGGCGTACTGACGGA TTCAT CGTTGGGGTCGGT 3′ and 98 nt: 5′ACCGAC CCCAACGATGAATCCGTCAGTACGCCAG TCGCC ACTGCCGGAGCCTTCATAAG CAATA TCAACAACGACGGACGCTGCCGGATTTACCTGCA3′.

Oligonucleotides for filaments interacting with 180 bp dsDNA. 82(−8) plus 16 heterologous for N= 82: 5′GACGCTG CCATATT CAAGTCGCCACTGCCGGAGCCTTCATAAGCAATATCAACAACG ACGGACGCTGCCGGATTTA CCTGCACTCCACGCCACTGAGG 3′; 50(−8) plus 48 heterologous for N= 50 5′ACGCTGCCATATTCAATCGTTCACTTTATTGCTGGTGTCATTGCTTGCTCA ACAACGACG GACGCTGCCGGATTTACCTGCACTCCACGCCACTG AGG 3′; 20(−8) plus 78 heterologous for N= 20: 5′GGACGCTGCCTTATTCCTGTTGAGTTTATTGCTGCCGTCATT GCTTATTATGTTCATCCCGTCA ACATTCAAACTGTTTGCACTCCACGCCACTGAGG 3′; 5(−8) plus 93 heterologous for N= 5: 5′GGACGCTGCCTTATTCCT GTTGAGTTTATTGCTGCCGT CATTGCTTATTATGTTCATCCCGTCAACATTCAAACTGTTCAGGGACGAATATGGTGAGG 3′; 8 heterologous 82 homologous plus 8 heterologous: 5′GACATTATAGTACGCCAGTCGCCACTGC CGGAGCCTTCAT AAGCAATAT CAACAACGACGG ACGCTG CCGGATTTACCTGCACTCC ACGCGCTGCCAT 3′; 42(−8) 56 heterologous for N= 42: 5′GGACGCTGCCTTATTCCTGTTGAG TTTATTGCTGCCGTCATTGCTTATTATGTTCAACT CCCGGGTATATGAAAGAGACGA CCACTGCCAGGGACGAA 3′; 98 nt Heterologous: 5′CGGAAAAGTGCATATCCAGCAGAA CATCATGAAAATAATGGGTAC T GTAAAAGC GGTGC CAGTCGGCATACTCCGTGGAT GACA TCCCGGCAAGCATG 3′.

Oligonucleotides annealed for 98 bp dsDNA and end labels: 5′/56-TAMN/CGGAAATCACTCCC GGGTATATGAA AGAGACGACCACTGCCAGGGACGAAAGTGCAATGCGGCATACCT CAGTGGCGTGGAG TGCAGGTATACAGATT 3′ and 5′AATCTGTATACCTGCACTCCACGCCA CTGAGGTATGCCGCATTGCACTTTCGTCCCTGGCAGTGGTCGTCTCTTTCATATACCCGG GAGTGATTTCCG/36-FAM/ 3′.

Oligonucleotides for filaments interacting with 98 bp dsDNA. 83 (−15) plus 15 heterologous: 5′ GGACGCTGCCGGA TTAATCTGTATACCTGCACTCCACGCCACTGAGG TATGCCGCATTGCA CT TTCGTCCCTGGCAGTGGTCGTCTCTTTCATATACC 3′; 75 (−15) plus 23 heterologous: 5′ GGACACTGCTTCATTCCTCTTATTACCTGCACTCCACG CCACTGAGGTAT GCCGCATTG CACTTTCGTCCCTGGCAGTGGTCGTCTCTTTCATATACC 3′; 50 (−15) plus 48 heterologous: 5′ GGACGCTGCC GGATTCCTGTTGAGTTTATTGCTGC CGTCATTGCTTATATGCCGCATTGCA CTTTCGTCCCTGGCAGTGGTCGTCTCTTTCATATACC 3′; 36 (−15) plus 62 heterologous: 5′ GGA CGCTGCCGGATTCCTGTTGAGTTTATTGCTGCCGTC ATTGC TTATTATGTTCATC CCGTTT TCGTCCCTGGCAGTGGTCGTCTCTTTCATATACC 3′; 20 (−15) plus 78 heterologous: 5′ GGACGC TGCCGGATTCCTGTTGAGTTTATTGCTG CCGTCATTGCTTATTATGTTCATCCCGTCAACAT TCAAACGGCCGGTCGTCTCTTTCATATACC 3′

### Analysis of genomes for repeated sequences and Chi sites

#### Genomes used

*Escherichia coli* O157, *E. coli* O157 strain 644-PT8, *E. coli* strain RR1, *E. coli* O157:H7 strain FRIK2533, *Salmonella enterica* subsp. *arizonae* serovar 62:z4,z23:-strain RSK2980, *Salmonella enterica* subsp. *enterica serovar Anatum* str. CDC 06-0532 strain USDA-ARS-USMARC-1764, *Shigella boydii* strain ATCC 9210, *S. flexneri* 5 str. 8401, *Klebsiella pneumoniae* subsp. *pneumoniae* strain TGH8, *K. pneumoniae* subsp. *pneumoniae* strain TGH10, *Proteus mirabilis BB2000,* and *Proteus mirabilis strain AR_005.*

For each of the genomes, the given strand is the strand given by the database from which we obtained the sequence. The sequences for the given strands of DNA for *E. coli* genomes were acquired from PATRIC in FASTA format. They were converted to a simple .txt file with A, C, G, and T bases and read into Matlab as a single continuous string running from 5′ to 3′ called *bases*. The sequence of the comp strand is the complement of the given strand; however, if each base in the .txt file for the given strand is simply replaced by the complementary base, the resulting comp strand sequence runs from the 3′ end to the 5′ end. To get the comp strand sequence running from 5′ to 3′, the order of the bases in the comp strand must be reversed.

#### Repeated sequences in whole genomes

To find all repeated sequences within the whole genome, 20 bp was established as an important cutoff length, and all the starting positions in which each consecutive sequence of 20 bp occurred were mapped within the genome. Sequences and their starting locations that were repeated were selected and placed in a smaller map “*g_rep*”. Due to the overlap of these 20 bp keys, repeated sequences longer than 20 bp would register more than one key within “*g_rep*”. In order to determine the true starting positions of repeated sequences, the multiple starting positions associated to a particular 20 bp sequence were retrieved, but isolated from groupings of starting positions of other 20 bps sequences. A comparison list *“complist”* was generated to choose all the comparisons within each group. For a 20 bp sequence with only two starting positions, there was only one comparison. But for sequences with *n* starting positions, there were *C*(*n*,2) (n choose 2) comparisons to be made. All comparisons were made against an arbitrarily large genome section of 10,000-20,000 bp on either side of the starting position for the two sequences being compared. The first mismatch in either direction was found and its distance to the starting position as well as its absolute location in “*bases*” was recorded. If there were conflicts between two comparisons within the same group, indicating that at least one sequence in the group was a subsequence of the others, the maximum distance was chosen only for the sequences where the conflict occurred. Therefore, not all sequences within a particular grouping necessarily have the same distance.

In the resulting array of start and end position pairs, all repeats were discarded, as these are a remnant of the over-counting from the original selection of positions. The unique start and end positions represent the starting and ending positions of all sequences >= 20 bp in the whole genome that occur more than once. From this, the length of homology is easily calculated for each particular sequence, and start, difference, and end information was succinctly summarized in array “ *start_difference_end”*.

#### Chi sites

Using MatLab’s built in function to find the location of substrings, the starting indexes of all the Chi sites (5′-GCTGGTGG-3′) (Smith et al., 1981) on the given strand going from 5′ to 3′ were found. Similarly, positions for the reverse complements of the Chi sites were also found to represent the position of Chi sites on the complementary strand, which was not read into MatLab and therefore not directly searched.

#### Probability mass function for L_chi_

For each position in “*bases*”, the distance to the nearest Chi sites in both directions was calculated using a “*loopindex*” function. The distances were summed for each position. Using MatLab’s default “*histogram*” function with bin sizes of 5,000 bp, the results were acquired for each *E. coli* genome. The bin counts for each were averaged and normalized to represent the case in which the position of the DSB is assumed to be random and RecBCD is assumed to recognize Chi sites with 100% accuracy. For the case where RecBCD only has an ~ 30 % chance of recognizing a Chi site, the DSB position was still assumed to be random, but the number of Chi sites skipped for each break was generated using a first success distribution in MatLab:

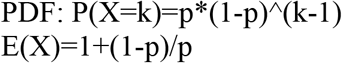

where X is the random variable denoting the number of Chi sites up to and including the recognized Chi site, p is the probability that a particular chi site is recognized, and E(X) is the expectation value for the random variable with a given p. Adjusted distances for each position were then calculated and a new histogram with bin size 10,000 was generated for each *E. coli* genome. The individual bin counts were averaged for the four genomes and normalized.

#### Repeats adjacent to Chi sites

A method similar to the one used for N_repeat_ was used to find repeats >= 20 bp adjacent to a Chi site that would remain as part of the searching filament (N_rep_ 3′). For Chi sites on the given strand, the 20 bp to the 5′ end of the start location of the Chi site in “*bases*” was selected as the key; for Chi sites on the comp strand, the complement of the 20 bp to the 3′ end of the comp Chi site on the given strand was selected as the key. Those are the 20 bp on the 5′ side of the Chi site on the comp strand. For each type of Chi, a particular sequence key was mapped to the starting position(s) of the associated Chi site(s). We did not consider interactions between sequences in the given strand and sequences in the complementary strand. Thus, the four possible interactions *xgiven_pos, xgiven_rep, xcomp_pos,* and *xcomp_rep* mapped unique and repeated sequences to their associated Chi sites for each Chi type.

The actual lengths of the repeats were found in a way similar to the lengths of repeats found in the entire genome. The result was a table of starting positions of Chi sites with N_rep 3′_ >= 20 bp and the actual length of homology to either side of the starting position of the Chi site.

#### Distances between repeat adjacent and nearest Chi sites

For each Chi site whose N_rep 3′_ >= 20, the distance to the nearest Chi site of the same type in the direction of strand exchange progression was found. The next Chi site in the sorted list was selected, and its difference was calculated. This distance represents the number of positions where, if a DSB were to occur, that Chi site would be first encountered by RecBCD in the RecBCD pathway. Dividing this number by the number of bp in the genome gives the fraction of the genome that would result in that particular searcher if DSB occurred randomly and RecBCD was 100 % accurate in identifying a Chi site.

#### Fraction of genome that gives Chi and WG searcher

Selecting for repeat length greater than or equal to *N = [0,100,200 … 16000]*, the fractions were found and summed over all Chi sites of one type as well as overall. The results were displayed using MatLab’s *plot* function. Similar fractions were calculated for whole genome (WG) repeats where one or both sides of the remaining sequence are required to be N. Sequences of at least N were found. Subtracting each by N, multiplying by 2, and summing together gave the raw number of positions that resulted in at least one side having the requisite number of homology. Taking the same sequence of at least N, subtracting 2*N from each (choosing the max of the result or 0), and summing over all gives the raw number of positions that results in both sides. Dividing the raw number by the number of bases gives the fraction.

**Table 1.** Abbreviations used in the text

| Abbreviation | Meaning |
| --- | --- |
| 3p end | The end of the dsDNA that would be reached by synthesis initiated by RecA mediated recombination at the 3' end of the initiating ssDNA |
| $\Delta F$ | The change in fluorescence with time |
| $\Delta\Delta F$ | The difference between $\Delta F$ for the positive and the $\Delta F$ for a control with $N=20$ |
| $\Delta L$ | The separation between the fluorescent labels and the 3' end of the initiating ssDNA |
| $D_{\text{label}}$ | The separation between the fluorescent labels and the 3p end of the dsDNA |
| $D_{\text{init}}$ | The separation between the 3' end of the initiating ssDNA and the 3p end of the dsDNA |
| $\text{DSB1}_{\text{frac}}(n)$ | The fraction of the DSBs that creates initiating strands with $N_{\text{rep } 3'} > n$ on a specified initiating strand. |
| $\text{DSB2}_{\text{frac}}(n)$ | The fraction of the DSBs that creates initiating strands with $N_{\text{rep } 3'} > n$ on both initiating strands |
| $L_{\text{Chi}}$ | The number of bases surrounding the DSB that are not incorporated in the searching filaments because they are removed by RecBCD. |
| $L_{\text{prod}}$ | The length of a heteroduplex product joining the initiating and complementary strands |
| $M_{3'}$ | The number of contiguous mismatched bp at the 3' end of the initiating ssDNA |
| $N$ | The number of contiguous bp in the dsDNA that are sequence matched to bases in the initiating ssDNA in experiments with only one initiating ssDNA |
| $N_1$ | The number of contiguous bp in the dsDNA that are sequence-matched to bases in one of the initiating ssDNA in experiments with two initiating ssDNAs |
| $N_2$ | The number of contiguous bp in the dsDNA that are sequence matched to bases in the other of the initiating ssDNA in experiments with two initiating ssDNA |
| $N_{\text{repeat}}$ | The length of a repeated sequence occurring anywhere in the genome |
| $N_{\text{rep } 3'}$ | The length of a repeated sequence that is positioned on the 5' side of a Chi site. In the RecBCD pathway, these repeats would occur at the 3' end of searching filaments. |

## Acknowledgements

T.F.T. and V.G. would like to thank members of the Godoy Lab for their help, especially Margaret Downs. M.P. would also like to acknowledge useful conversations with Prof. Phillip Sharp. The work was supported by Northeastern University and NIGMS RO1GM088230 to VG. C.P. was supported by ‘Initiative d’Excellence’ program of the French State (‘DYNAMO’, ANR-11-LABX-0011-01. Funding for C.L. was provided by HCRP (Harvard College). Support was also provided by Harvard University.

## Author contributions

C.L. performed almost all analysis of bacterial sequences; C.D. performed all experiments, and contributed to the experimental design and data analysis; T.F.T. and V.G provided the DNA Pol IV protein, contributed to the experimental design, and offered structural insight; C.P. contributed to the understanding of the interaction between Pol IV and RecA; and M.P. conceived the study, contributed to the analysis of bacterial sequences, experimental design, and data analysis. All of the authors discussed the results, contributed to the manuscript preparation, and approved the final version of the manuscript.

---

**Figure 3- figure supplement 1.**
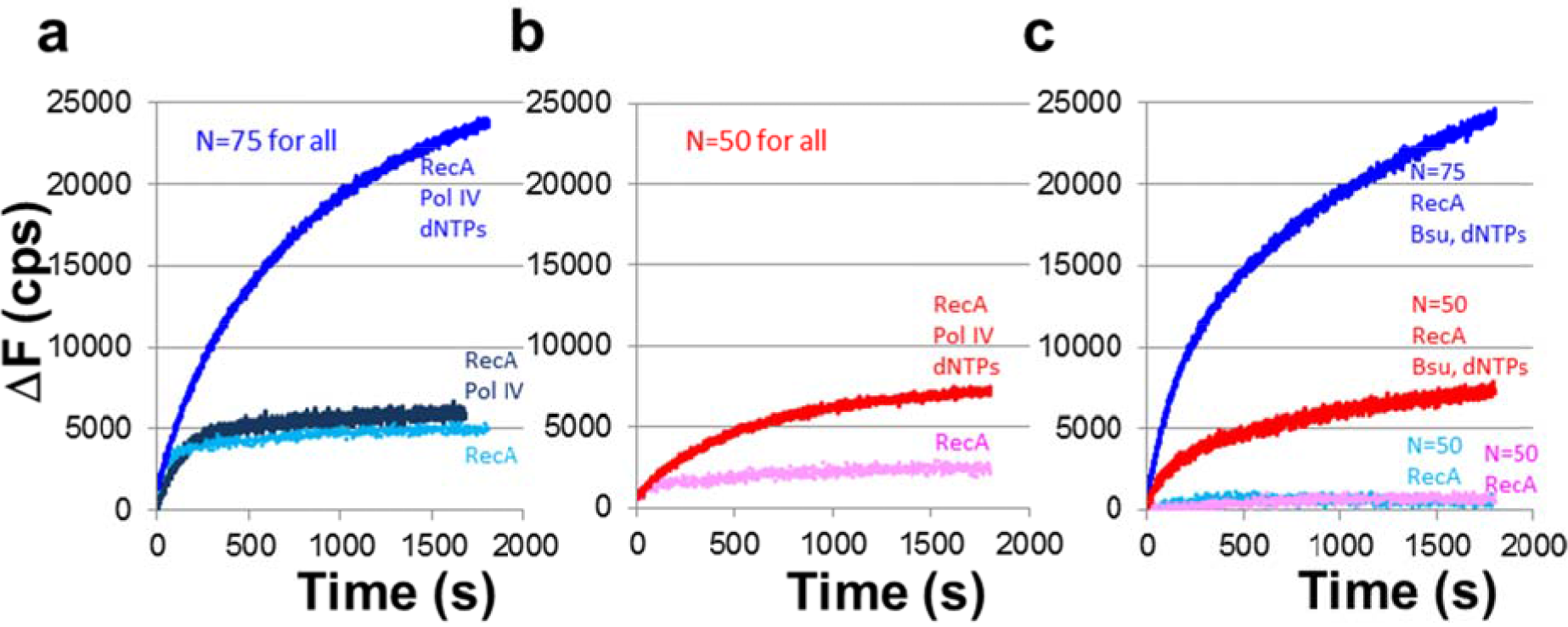
ΔF vs. time curves in the presence and absence of DNA polymerase where initial fluorescence values for the heterologous filament have been subtracted from the observed values. **a** ΔF vs. time curves for N = 75 in the presence of dATP-ssDNA-RecA filaments (RecA), DNA Pol IV, and dNTPs (royal blue); analogous results with dATP-ssDNA-RecA filaments and DNA Pol IV but without dNTPs (dark blue); results in the presence of dATP-ssDNA-RecA filaments only (light blue). **b** ΔF vs. time curves for N = 50 in the presence of dATP-ssDNA-RecA filaments, DNA Pol IV, and dNTPs (red), and dATP-ssDNA-RecA filaments only (pink). **c** ΔF vs. time curves in the presence of ATP-ssDNA-RecA filaments, LF-Bsu, and dNTPs for N = 75 (blue) and N = 50 (red), and results in the presence of ATP-ssDNA-RecA filaments only for N = 75 (light blue) and N = 50 (pink).

**Figure 3- figure supplement 2.**
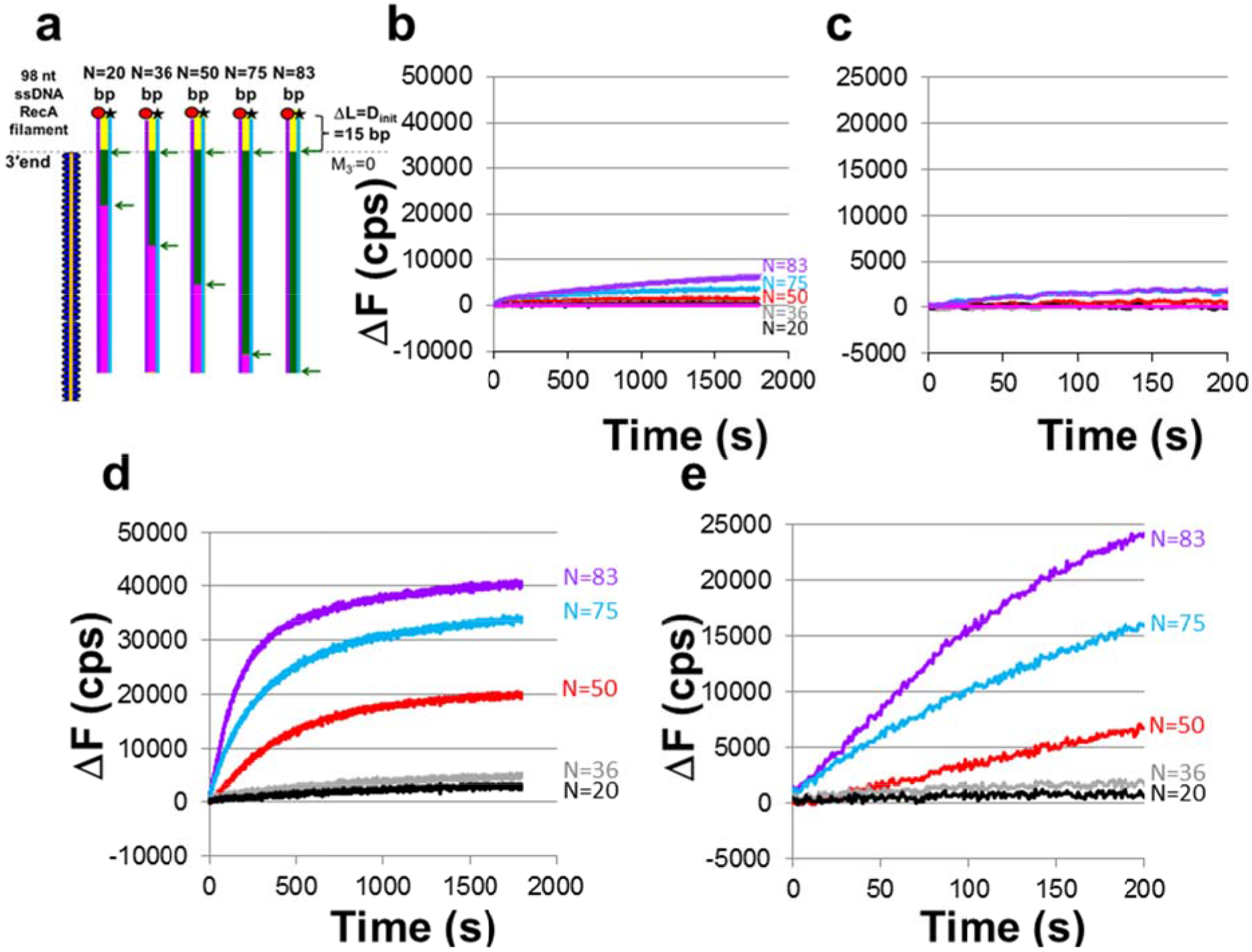
ΔF vs. time curves in the absence and presence of LF-Bsu polymerase. **a** Schematic of the experiments. Each of the ssDNA-RecA filaments was formed on a 98 nt ssDNA. One such filament is illustrated by the orange line with blue ellipses. Each representation of the same 98 bp dsDNA is colored to indicate the region of the dsDNA containing the N contiguous bases that are sequence matched to the bases in each of the different initiating strands. The N matched bases are shown in green and highlighted by green arrows. The N value for each illustration is listed above the dsDNA. Non-sequence matched bases within and beyond the matching filament region are shown in magenta and yellow, respectively; N values are 20, 36, 50, 75 and 83. **b** iΔF vs. time curves in the presence of ATP-ssDNA-RecA filaments without LF-Bsu for N = 83 (purple), 75 (blue), 50 (red), 36 (gray), 20 (black), and heterologous ssDNA-RecA filament (magenta). **c** First 200 s of data without polymerase shown in (**b**). **d** ΔF vs. time curves in the presence of ATP-ssDNA-RecA filaments and LF-Bsu where the heterologous curve was subtracted for N = 83 (purple), 75 (blue), 50 (red), 36 (gray), and 20 (black). **e** First 200 s of data with LF-Bsu shown in (d).

**Figure 4- figure supplement 1.**
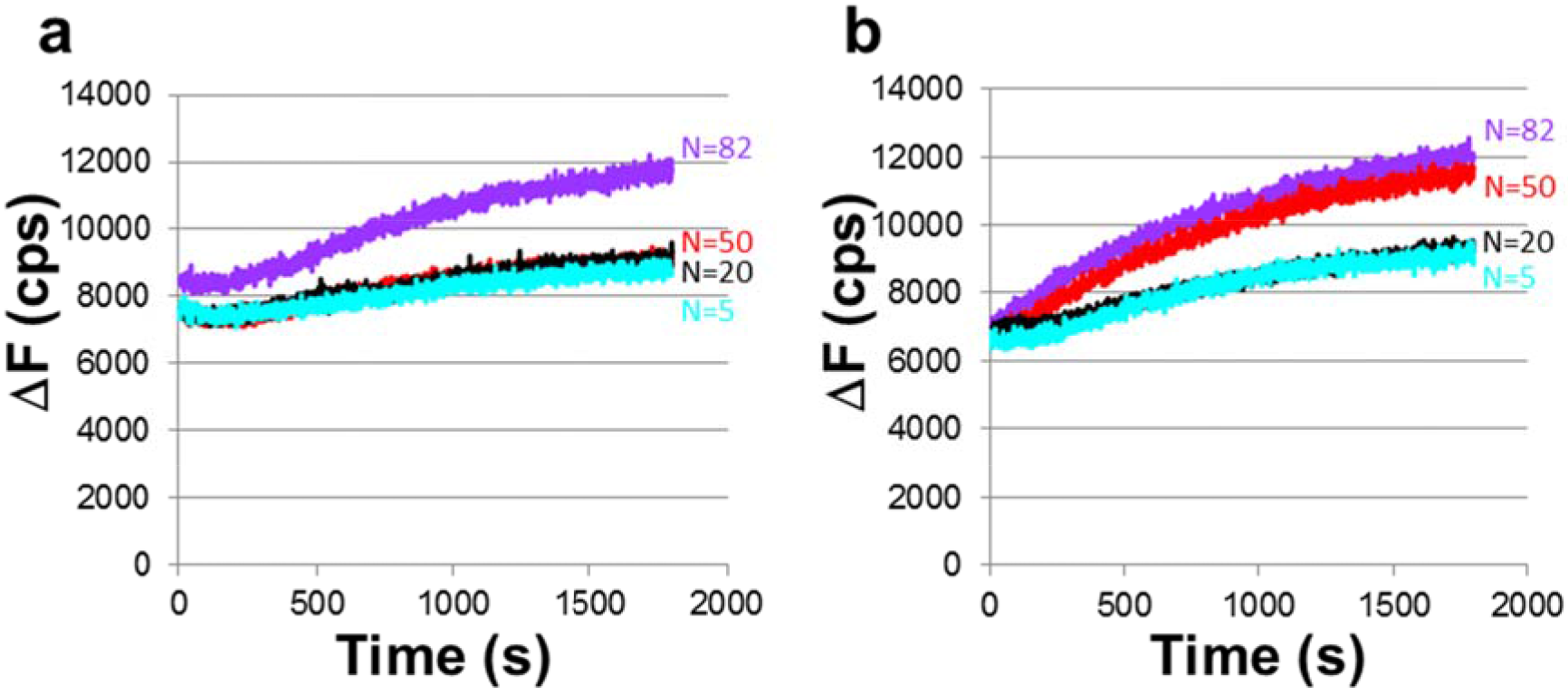
Raw fluorescence vs. time curves for dATP-ssDNA-RecA filaments, DNA Pol IV, and 180 bp labeled dsDNA construct; data shown in Figure 4b, d. **a** ΔF vs. time curves for raw data corresponding to Figure 4b and N = 82 (purple), 50 (red), 20 (black), and 5 (cyan). **b** ΔF vs. time curves for raw data corresponding to Figure 4d and N = 82 (purple), 50 (red), 20 (black), and 5 (cyan).

**Figure 4- figure supplement 2.**
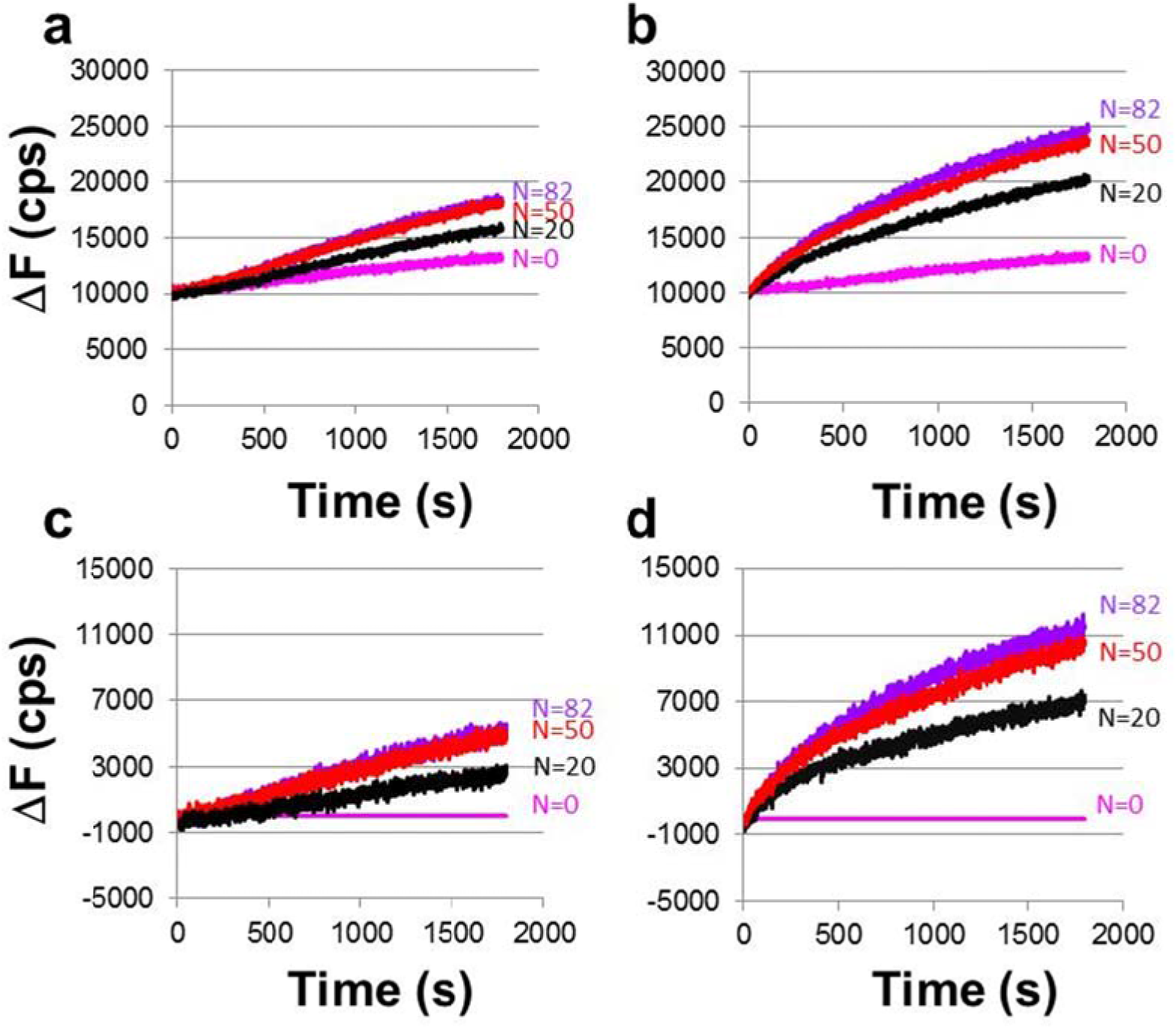
ΔF vs. time curves obtained with ATP-ssDNA-RecA filaments, LF-Bsu polymerase, and 180 bp labeled dsDNA construct. The purple, red, black, and magenta curves correspond to N = 82, 50, and 20, and heterologous DNA, respectively. a Raw data for one filament experiments with LF-Bsu, represented by the schematic shown in Figure 4a. **b** Raw data for two filament experiments with LF-Bsu, represented by the schematic shown in Figure 4c. **c** ΔF vs. time curves shown in (a) after subtracting the heterologous DNA curve. d AF vs. time curves shown in (b) after subtracting the heterologous DNA curve.

**Figure 5- figure supplement 1.**
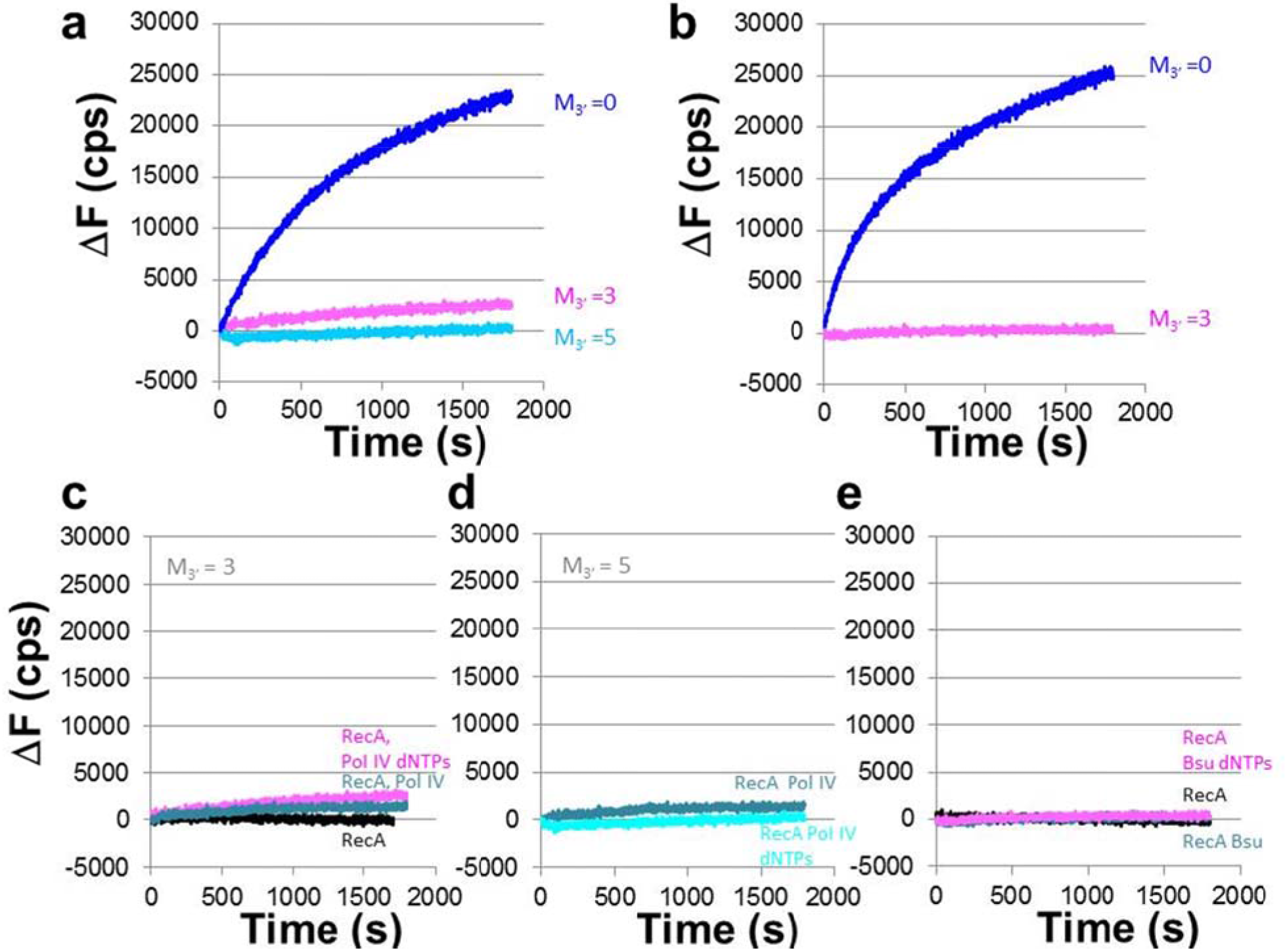
Control experiments for the data shown in Figure 5b, c. **a** Same as Figure 5b; ΔF vs. time curves in the presence of dATP-ssDNA-RecA filaments (RecA), DNA Pol IV, and dNTPs where the blue, light blue, and pink curves correspond to M_3′_ values of 0, 3, and 5 base mismatches, respectively. **b** Same as Figure 5c; results in the presence of ATP-ssDNA-RecA filaments, LF-Bsu polymerase, and dNTPs. **c** Interactions with M_3_ =3 dATP-ssDNA-RecA filaments without any DNA polymerase (black), with M_3′_ =3 dATP-ssDNA-RecA filaments, DNA Pol IV, and no dNTPs (blue), and with M_3′_ =3 dATP-ssDNA-RecA filaments, DNA Pol IV, and dNTPs (pink). d Results with M_3′_ = 5 ssDNA-RecA filaments, DNA Pol IV, and no dNTPs (blue-green) and with M3′ = 5 ssDNA-RecA filaments, DNA Pol IV, and dNTPs (cyan). **e** Results are analogous to (**c**) with =3 ATP-ssDNA-RecA filaments and LF-Bsu.

**Figure 5- figure supplement 2.**
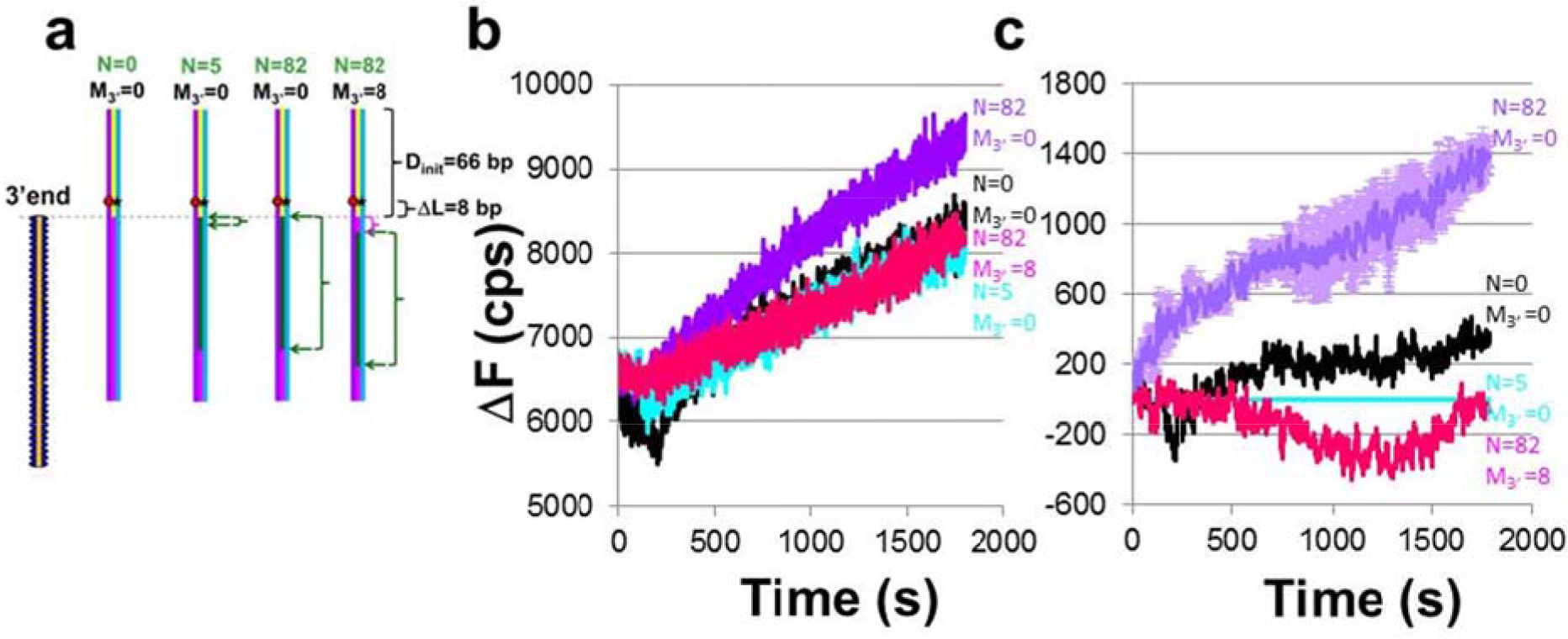
2M3′ dependence of DNA synthesis by DNA Pol IV. **a** Schematic of the expermient. The orange, purple, andblue lines indicate the initiating, complementary and outgoing stands respectively The blue ovals indicate the RecA in the initial ssDNA-RecA filament. Different ssDNA sequences are used presenting regions of homology of variable length with respect to the 180 bp dsDNA construct. The green rectangles indicate regions of the dsDNA target that are sequence matched to the corresponding bases in the initiating strand. The magenta rectangle indicates regions of the dsDNA that are heterologous to the corresponding bases in the initiating strand. The dashed line indicates the position of the 3′ end of the ssDNA-RecA filament when the green region is adjacent to the sequence matched bases in the ssDNA-RecA filament. The yellow region indicates bases in the dsDNA that are beyond the 3′ end of the ssDNA-RecA filament and heterologous to the bases in the filament. **b** ΔF vs. time curves for dATP-ssDNA-RecA filaments and DNA Pol IV for N = 82 M_3_= 0 (purple) and N = 82 M_3_, = 8 (dark pink). Controls with no mismatches (M_3_, = 0) are also shown by the cyan (N = 5) and the black (N = 0) curves. **c** ΔΔF vs. time curves where raw fluorescence vs. time for N = 5 in (**b**) was subtracted from the other raw fluorescence vs. time curves. The light purple error bars correspond to the standard deviation for two repetitions of the N = 82 M_3_, = 0 data.

**Figure 6- figure supplement 1.**
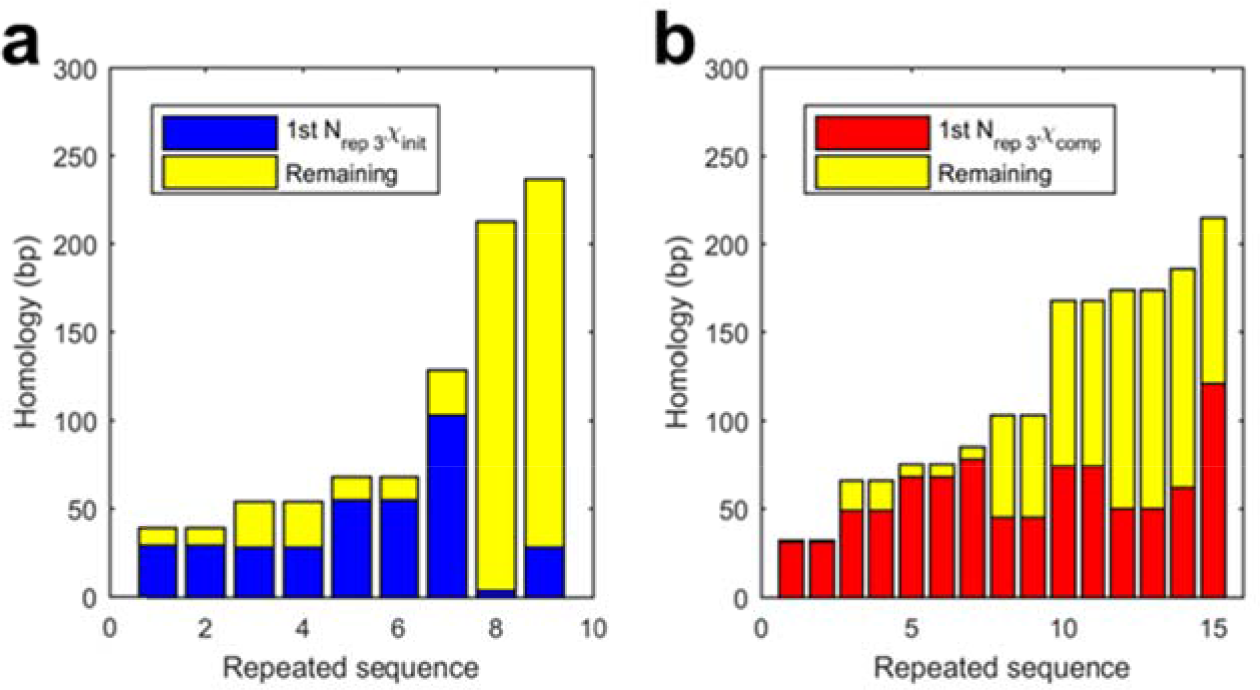
Expanded view of Figure 6a, b shown in the main text. **a** Repeats in the given strand that include a Chi site. The total height of the bar corresponds to the total length of the repeat. The height of dark blue regions corresponds to the part of the repeat that is on the 5′ side of the Chi site that is most distant from the 3′ end of the repeat. The height of the yellow regions corresponds to the portions of the long repeat that would be degraded by RecBCD because they are on the 3′ side of both the Chi sites; x-axis expanded to highlight the 9 shortest repeats. **b** Analogous results for the comp strand. The regions that are colored dark blue in (a) are colored red in (b).

**Supplementary Table 1.** Comparison between occurrences of Chi sites within repeats longer than 20 bp and analogous occurrences for an equal number of randomly positioned markers. Separate results are shown for each of 12 enteric bacteria.

|  | ecoli_MP |  |  | ecoli1 |  |  | ecoli2 |  |  |
| --- | --- | --- | --- | --- | --- | --- | --- | --- | --- |
|  | random | st dev | actual | random | st dev | actual | random | st dev | actual |
| # given strand hits | 40.55 | 6.93 | 23 | 59.39 | 7.74 | 32 | 16.41 | 3.81 | 6 |
| # comp strand hits | 39.75 | 5.65 | 27 | 61.26 | 7.92 | 35 | 16.65 | 3.86 | 3 |
| # >=2 given same repeat | 3.45 | 1.70 | 5 | 6.54 | 1.87 | 3 | 1.76 | 0.93 | 1 |
| # >=2 comp same repeat | 3.58 | 1.64 | 3 | 6.97 | 1.89 | 5 | 1.74 | 0.85 | 1 |
| # of repeats containing hits on each strand.No Chi site pair is correctly positioned to create filament pair that includes the same repeat. Incorrectly positioned Chi site pairs are highlighted in yellow | 4.71 | 1.94 | 0 | 9.61 | 2.16 | 0 | 1.67 | 1.16 | 0 |
| Number of Chi sites in comp strand |  |  | 551 |  |  | 595 |  |  | 506 |
| Number of Chi sites in given strand |  |  | 571 |  |  | 581 |  |  | 493 |
|  | ecoli5 |  |  | salmonella1 |  |  | salmonella2 |  |  |
|  | random | st dev | actual | random | st dev | actual | random | st dev | actual |
| # given strand hits | 52.56 | 6.99 | 28 | 9.58 | 3.37 | 7 | 8.50 | 2.88 | 3 |
| # comp strand hits | 52.21 | 6.98 | 28 | 8.64 | 2.83 | 2 | 8.42 | 3.25 | 10 |
| # >=2 given same repeat | 5.94 | 2.20 | 5 | 1.33 | 0.72 | 1 | 1.30 | 0.55 | 1 |
| # >=2 comp same repeat | 5.85 | 2.00 | 3 | 1.22 | 0.43 | 1 | 1.34 | 0.58 | 2 |
| # of repeats containing hits on each strand.No Chi site pair is correctly positioned to create filament pair that includes the same repeat. Incorrectly positioned Chi site pairs are highlighted in yellow | 8.47 | 2.58 | 0 | 0.77 | 0.77 | 2 | 0.86 | 1.00 | 2 |
| Number of Chi sites in comp strand |  |  | 572 |  |  | 307 |  |  | 414 |
| Number of Chi sites in given strand |  |  | 578 |  |  | 349 |  |  | 415 |

Supplementary Table 1 continuation
|  | shigella1 |  |  | shigella2 |  |  | klebsiella1 |  |  |
| --- | --- | --- | --- | --- | --- | --- | --- | --- | --- |
|  | random | st dev | actual | random | st dev | actual | random | st dev | actual |
| # given strand hits | 53.47 | 6.83 | 11 | 41.25 | 6.06 | 8 | 21.24 | 4.28 | 16 |
| # comp strand hits | 52.33 | 6.34 | 2 | 44.29 | 5.65 | 11 | 20.24 | 4.61 | 12 |
| # >=2 given same repeat | 4.89 | 2.03 | 3 | 3.66 | 1.62 | 3 | 2.35 | 1.08 | 1 |
| # >=2 comp same repeat | 4.63 | 1.78 | 1 | 3.95 | 1.54 | 3 | 2.26 | 1.00 | 1 |
| # of repeats containing hits on each strand.No Chi site pair is correctly positioned to create filament pair that includes the same repeat. Incorrectly positioned Chi site pairs are highlighted in yellow | 7.15 | 2.60 | 0 | 5.20 | 2.43 | 0 | 2.86 | 1.38 | 0 |
| Number of Chi sites in comp strand |  |  | 463 |  |  | 499 |  |  | 924 |
| Number of Chi sites in given strand |  |  | 466 |  |  | 469 |  |  | 942 |
|  | klebsiella2 |  |  | proteus1 |  |  | proteus2 |  |  |
|  | random | st dev | actual | random | st dev | actual | random | st dev | actual |
| # given strand hits | 21.64 | 4.25 | 16 | 3.66 | 2.09 | 1 | 6.59 | 2.38 | 2 |
| # comp strand hits | 21.44 | 4.70 | 12 | 3.98 | 1.97 | 1 | 6.23 | 2.58 | 3 |
| # >=2 given same repeat | 2.27 | 1.19 | 1 | 0.91 | 0.40 | 0 | 1.23 | 0.49 | 1 |
| # >=2 comp same repeat | 2.27 | 1.05 | 1 | 0.93 | 0.28 | 0 | 1.17 | 0.52 | 1 |
| # of repeats containing hits on each strand.No Chi site pair is correctly positioned to create filament pair that includes the same repeat. Incorrectly positioned Chi site pairs are highlighted in yellow | 2.53 | 1.48 | 0 | 0.09 | 0.31 | 0 | 1.01 | 0.91 | 2 |
| Number of Chi sites in comp strand |  |  | 927 |  |  | 159 |  |  | 144 |
| Number of Chi sites in given strand |  |  | 941 |  |  | 146 |  |  | 161 |

Supplementary Table 1 continuation
| Sum over all Genomes |  |  |  |
| --- | --- | --- | --- |
|  | random | st dev | actual |
| # given strand hits | 334.84 | 17.91 | 153 |
| # comp strand hits | 335.44 | 17.40 | 146 |
| # both in same repeat<br>correctly oriented | 22.47 | 2.99 | 0 |
| # >=2 given same repea | 35.63 | 4.76 | 25 |
| # >=2 comp same repea | 35.91 | 4.41 | 22 |
| Number of Chi sites in comp strand |  |  | 6061 |
| Number of Chi sites in given strand |  |  | 6112 |

